# Variation in a pleiotropic hub gene drives morphological evolution: Insights from interspecific differences in head shape and eye size in *Drosophila*

**DOI:** 10.1101/2020.01.24.918011

**Authors:** Elisa Buchberger, Anıl Bilen, Sanem Ayaz, David Salamanca, Cristina Matas de las Heras, Armin Niksic, Isabel Almudi, Montserrat Torres-Oliva, Fernando Casares, Nico Posnien

**Author notes:** **Author contribution** EB: Conceptualization, Data curation, Formal analysis, Investigation, Methodology, Visualization, Writing – original draft, Writing-review and editing. AB: Investigation. SA: Investigation. DS: Investigation. CMH: Investigation. AN: Investigation. IA: Resources, Writing – review and editing, MTO: Investigation, Resources, Writing – review and editing, FC: Investigation, Writing-review and editing. NP: Conceptualization, Funding acquisition, Project administration, Resources, Supervision, Visualization, Writing – original draft, Writing – review and editing.

## Abstract

Revealing the mechanisms underlying the breath-taking morphological diversity observed in nature is a major challenge in Biology. It has been established that recurrent mutations in hotspot genes cause the repeated evolution of rather simple morphological traits, such as body pigmentation or the gain and loss of structures. To date, however, it remains elusive whether hotspot genes contribute to natural variation in complex morphological traits, such as the size and shape of organs. Since natural variation in head morphology is pervasive in *Drosophila*, we studied the molecular and developmental basis of differences in compound eye size and head shape in two closely related *Drosophila* species. We show that differences in both traits are established late during head development and we applied comparative transcriptomics and chromatin accessibility data to identify the GATA transcription factor Pannier (Pnr) as central factor regulating these differences. Although the genetic manipulation of Pnr affected multiple aspects of dorsal head development, the effect of natural variation is restricted to a subset of the phenotypic space. We present data suggesting that this developmental constraint is caused by the co-evolution of expression of *pnr* and its co-factor *u-shaped (ush)*. We propose that natural variation in highly connected developmental regulators with pleiotropic functions is a major driver for morphological evolution and we discuss implications on gene regulatory network evolution. In comparison to previous findings, our data strongly suggests that evolutionary hotspots do not contribute to the repeated evolution of eye size and head shape in *Drosophila*.

## Introduction

The morphological diversity present in nature is a key prerequisite for organisms to adapt to an ever-changing environment. Since the genome of an organism contains instructive information about its morphology, one of the major goals in biological research is to establish genotype-phenotype correlations for a given morphological trait [1,2]. The genetic architecture of relatively simple traits has been successfully determined at high resolution. For instance, natural variation in body pigmentation in the vinegar fly *Drosophila melanogaster* and the beach mouse *Peromyscus polionotus* has been mapped to individual nucleotides affecting gene expression [3,4] or protein function [5], respectively. Also, the genetic basis of gain or loss of structures like trichomes in *Drosophila* [6], armor plates in three spine stickleback fish [7] or the repeated loss of eyes in cave fish [8,9] has been successfully revealed. Intriguingly, the analysis of similar morphological differences in various lineages showed that the same genes were repeatedly affected. For instance, differences in abdominal pigmentation and the occurrence of wing spots across the phylogeny of Drosophilidae have been mapped to the *bric-à-brac* (*bab*) [10] and *yellow* (*y*) [11] loci, respectively. Also the evolution of larval trichome patterns in various *Drosophila* species is caused by variation in the regulation of the *shavenbaby* (*svb*) gene [12] and recurrent natural variation in the *Pitx1* locus has been causally linked to pelvic spine reduction in different stickleback lineages [13,14]. While recurrent natural variation in such hotspot genes seems to be common for the evolution of rather simple morphological traits, it remains elusive, whether evolutionary hotspots play a major role in the evolution of complex traits, such as the size and shape of organs.

The identification of the molecular changes underlying variation in complex morphological traits is difficult because many genomic loci with small effect sizes that are spread throughout the genome contribute to trait differences [15–17]. For instance, natural variation in mandible and craniofacial shape in mouse is influenced by various loci located on most of the chromosomes [18–21] and studies in *Drosophila* revealed loci on several chromosomes associated with differences in eye size and head shape [22–24]. The omnigenic model suggests that most genomic loci can influence the phenotypic outcome even though they are not functionally linked to the trait (“peripheral genes”) because they are connected to loci with a direct effect (“core genes”) in highly wired gene regulatory networks (GRNs) [15]. As many other biological networks [25–28], GRNs are thought to be scale-free, meaning that most nodes within the network have only very few connections and these low-degree nodes are connected by a few highly connected nodes, so called hubs [29,30]. Genes that act as hubs relay complex incoming information to a high number of target genes and they often code for evolutionary conserved proteins with crucial functions within the network [31,32]. Therefore, it is conceivable that “core genes” proposed by the omnigenic model have features of hub genes within GRNs. It has been suggested that the recurrent evolution in hotspot genes is facilitated by their hub position within GRNs [12,33,34], as well as the observation that only few such hubs are necessary to orchestrate the formation of rather simple traits [35]. In contrast, the development of complex organs relies on the interplay of many hub genes [36,37], suggesting that evolutionary hotspots are less likely to be relevant for complex trait evolution.

An excellent model to test, whether natural variation in complex traits is caused by recurrent changes in hotspot genes is the *Drosophila* head that harbours major sensory organs, such as the compound eyes [38]. Natural intra- and interspecific variation in eye size is pervasive in various *Drosophila* lineages and increased eye size is often associated with a reduction of the interstitial cuticle between the eyes [23,39–42]. While the genetic architecture of this trade-off is starting to be revealed in various species [22–24,43], the most comprehensive genetic and developmental analysis established that intra- and interspecific differences in the trade-off are caused by variation in the early subdivision of the eye-antennal imaginal disc [44], the larval tissue from which most of the adult head develops [45]. Genetic analyses between different *D. melanogaster* strains revealed a single nucleotide polymorphism in the regulatory region of the *eyeless/pax6* (*ey*) gene affecting its temporal expression and thus eye-antennal disc subdivision [44]. Ey regulates various target genes [46] and signalling pathways [47] to initiate retinal development, suggesting that it acts as a hub gene during eye development. Additionally, the recurrent variation in *ey* expression suggests that *ey* might also be a hotspot gene regulating the trade-off between eye and head cuticle development [44]. We sought to test, whether variation in the early subdivision of the eye-antennal disc is a common mechanism underlying the development of a head trade-off in *Drosophila*. As model we studied head development in the two closely related *Drosophila* species *D. melanogaster* and *D. mauritiana*, which vary in adult eye size and head shape [23,42,48]. Specifically, *D. mauritiana* develops larger compound eyes with more ommatidia and the bigger eyes of *D. mauritiana* are accompanied by a narrower interstitial head cuticle [42].

In contrast to previous reports, we could show that differences in compartment sizes occur only late during eye-antennal disc development, suggesting that the early subdivision of the disc is not affected. To reveal candidate genes responsible for eye size differences, we developed a new method to identify evolutionary relevant hub genes with crucial positions within the developmental GRN. Applying comparative transcriptomics and functional genomics, we revealed the GATA transcription factor Pannier (Pnr) and its transcriptional co-factor U-shaped (Ush) as candidate hub genes. We functionally confirmed that the overexpression of *pnr* in *D. melanogaster* indeed phenocopies aspects of the *D. mauritiana* head shape as well as eye size. In summary, we argue that similar complex morphological differences can be caused by different developmental processes and molecular changes in different lineages. Therefore, our data suggests that evolutionary hotspots may play a less prominent role during complex trait evolution.

## Results

### Differences in dorsal head shape between *D. melanogaster* and *D. mauritiana* are defined during late eye-antennal disc development

*Drosophila* eye size and head shape vary extensively between *D. melanogaster* and *D. mauritiana* with the latter having bigger eyes due to more ommatidia at the expense of interstitial head cuticle [22,42,48]. Since eye size differences are most pronounced in the dorsal part [42], we first quantified differences in the dorsal head morphology to test whether dorsal head shape varies as well. We placed 57 landmarks on pictures of dorsal heads and applied a geometric morphometrics analysis (Fig 1A). A discriminate function analysis clearly distinguished the head shapes of *D. melanogaster* and *D. mauritiana* (Fig 1B) (p-value= 0.0001). In accordance with previous data [42], we found main differences in dorsal eye size with the eye area protruding more towards the back of the head in *D. mauritiana* (Fig 1B). The expansion of the eye area in *D. mauritiana* was accompanied by a narrower dorsal head region, which affected both the orbital cuticle and the dorsal frons region. The ocellar complex was slightly shifted ventrally in *D. mauritiana* (Fig 1B). Therefore, *D. melanogaster* and *D. mauritiana* do not only differ in the size of dorsal eye, but they also exhibit variation in the relative contribution of different head regions to the dorsal head.

**Fig 1.**
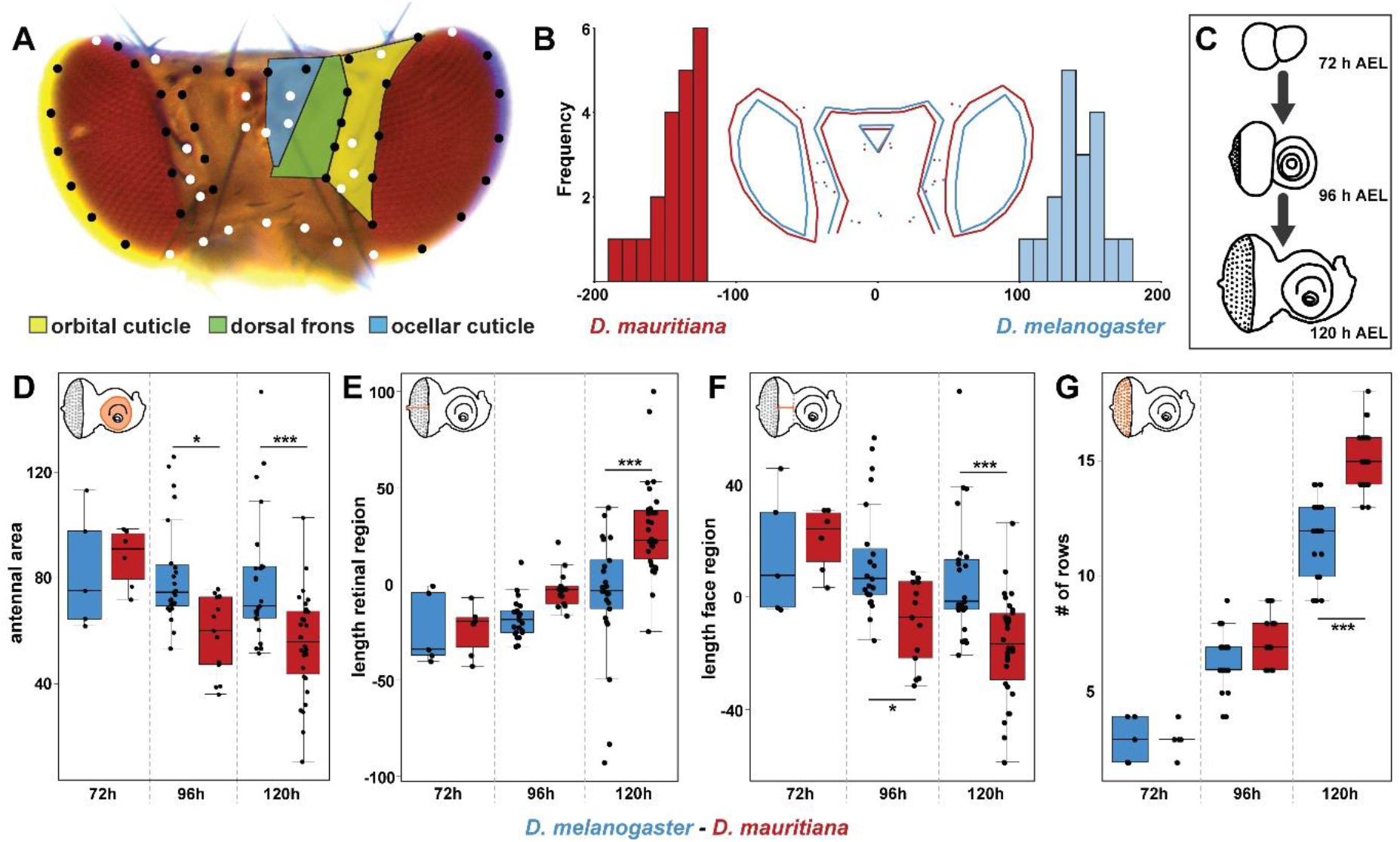
Quantitative differences in dorsal head shape are defined late during eye-antennal disc development. **(A)** Dorsal view of a *Drosophila melanogaster* head. The head cuticle consists of three morphologically distinguishable regions, namely the orbital cuticle (yellow) next to the compound eye, the dorsal frons (green) and the ocellary cuticle (blue). The 57 landmarks that were used to analyze head shape are shown as fixed (white) and sliding landmarks (black). **(B)** Discriminant function analysis distinguishes mean dorsal head shapes of *D. melanogaster* (blue) and *D. mauritiana* (red). Difference between means: Procrustes distance: 0.094, p-value= 0.0001 **(C)** The development was characterized and transcriptomic datasets were generated for developing eye-antennal discs for both species at three developmental stages: 72h after egg laying (AEL; late L2), 96h AEL (mid L3) and 120h AEL (late L3). **(D)** Area of the antennal region of the eye-antennal disc. Significant differences were observed at 96 h AEL and 120 h AEL (F5,96 = 7.86, p = 3.08e-6). **(E)** Distance from the optic stalk to the morphogenetic furrow was measured along the equator region. Significant differences were observed at 120 h AEL (F5,96 = 15.61, p = 3.2e-11). **(F)** Distance from the morphogenetic furrow to the antennal anlagen was measured. From 96 h AEL on, we observed significant differences (F5,96 = 10.23, p = 7e-8). **(G)** Number of ommatidial precursor rows was counted along the equator region of the eye-antennal disc. Significant differences were observed at 120 h after egg laying (AEL) (F5,96 = 210.8, p < 2e-16). One-way ANOVA followed by Tukey multiple comparisons: *** <0.001; * <0.05

To understand the developmental basis of the size and shape differences in dorsal head structures (Fig 1B), we compared eye-antennal disc development between species at the onset (72 h after egg laying; AEL), progression (96 h AEL) and termination of differentiation (120 h AEL) (Fig 1C). The size of the antennal and retinal region was not significantly different at 72h AEL, respectively (Fig 1D and 1E). First differences for both disc compartments were observed at 96 h AEL and differences were clearly visible at 120 h AEL. At these stages, *D. mauritiana* had a bigger retinal region and a smaller antennal region compared to *D. melanogaster* (Fig 1D and 1E). The region giving rise to the interstitial cuticle showed interspecific size differences from 96 h AEL on (Fig 1F) and the differences in the size of the retinal region was accompanied by significant differences in the number of ommatidia rows at 96 h and 120 h AEL (Fig 1G). Our data shows that the trade-off between head regions is defined late during eye-antennal disc development in *D. melanogaster* and *D. mauritiana*.

### Morphological differences between *D. melanogaster* and *D. mauritiana* are associated with variation in the developmental transcriptomic landscape

Next we tested whether differences in adult head morphology and eye-antennal disc development were associated with variation in the developmental transcriptomic landscape. We obtained comparative transcriptomes for eye-antennal disc development covering 72 h, 96 h and 120 h AEL (Fig 1C and S1 Fig) [49]. In line with the overall functional conservation of head morphology between species, we first confirmed that genes with stable interspecific expression represent central processes involved in eye-antennal disc development (S2 Fig). An analysis of those genes that were differentially expressed between species revealed that 72 % of variation in the dataset was due to differences between 72h and the other two stages (S3 Fig), an observation that was confirmed by a pairwise differential expression analysis. With 6,683 genes, we found the highest number of differentially expressed genes (DEGs) at 72 h AEL, while only 3,260 and 2,380 genes were differentially expressed at 96h AEL and 120 h AEL, respectively. This excess of DEGs at 72 h AEL is most likely caused by the fact that it is impossible to distinguish the sex of larvae at this early stage. Therefore, a mix of males and females was sequenced at 72 h AEL, while only females were analysed at the two later timepoints [49]. Differential expression was not biased towards one species since we observed an equal number of DEGs with higher expression in *D. melanogaster* and *D. mauritiana*, respectively (S1 Table).

To identify putative developmental processes affected by the interspecific expression differences, we clustered DEGs based on temporal expression profiles and then we searched for enriched GO categories associated with each gene cluster (Fig 2). Genes involved in generic metabolic and energy related processes were enriched in various clusters including those with dynamic (Fig 2, e.g. cluster 1 and 2) as well as stable temporal expression (Fig 2, e.g. cluster 11). Genes that tended to be highly expressed at later stages were enriched for factors regulating neuronal differentiation processes (Fig 2, e.g. cluster 4) and in clusters with stably expressed genes, we observed an enrichment for cell cycle control and tissue growth (Fig 2, e.g. clusters 8 and 14). In summary, we revealed extensive variation in the transcriptomic landscape of developing eye-antennal discs between *D. melanogaster* and *D. mauritiana* with different developmentally relevant processes being affected by these changes.

**Fig 2.**
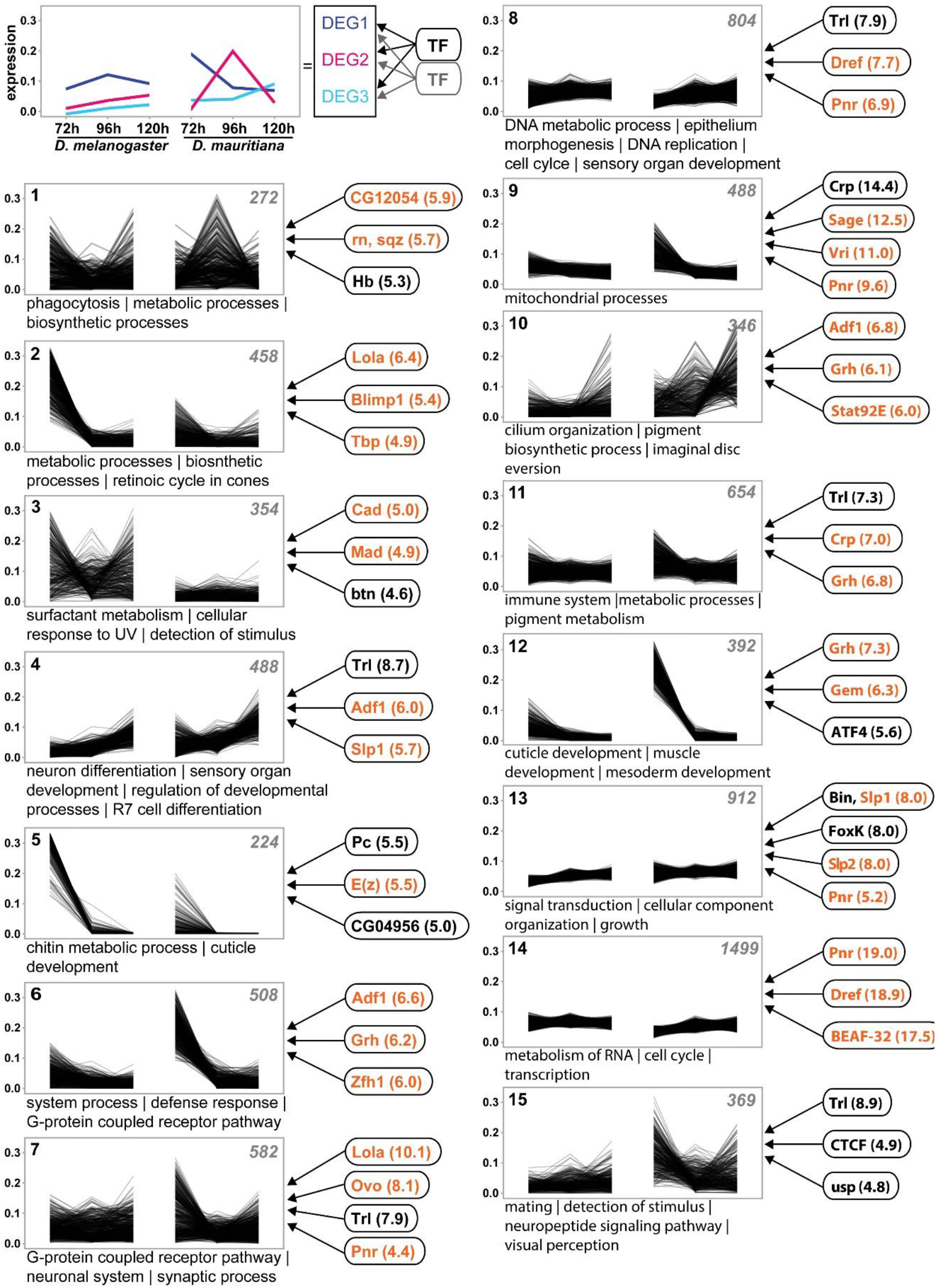
Extensive remodeling of the transcriptome underlying eye-antennal disc development. The scheme in the upper left corner illustrates the rational of the analysis. Differentially expressed genes (DEGs) with similar expression profiles were clustered and enriched transcription factor (TF) binding motifs were identified (see Materials and Methods for details). Cluster analysis of all differentially expressed genes between *D. melanogaster* and *D. mauritiana*. The number of genes in each cluster is given in the upper right corner. Significantly enriched GO terms are provided for each cluster. TFs refer to potential upstream regulators and the respective NES values (non-redundant occurrence and excluding non-*Drosophila* motifs). Factors written in orange are significantly differentially expressed in at least one stage between the two species. A complete list of enriched GO-terms and potential upstream factors is available in S2 Table.

### Variation in the transcriptomics landscape is caused by variation in expression of hub genes

The interspecific differences in gene expression dynamics must be the result of variation in the underlying regulatory interactions. Hub genes occupy central positions in developmental GRNs affecting many target genes [29,30]. To identify putative hub genes that regulate the expression of DEGs, we assume that DEGs with similar expression profiles are regulated by similar transcription factors; i.e. we treated co-expression clusters (clusters in Fig 2) as “gene modules” [50–52]. We established stage-specific chromatin accessibility data (ATACseq) to identify enriched transcription factor binding motifs for each cluster (see Materials and Methods and S1 Fig for details). Each cluster showed enrichment of motifs for a unique combination of transcription factors (Fig 2), suggesting that variation in different gene modules is influenced by different putative hub genes.

Many hub genes play central roles during development [31,32] and indeed, some of the identified transcription factors have previously been described to be involved in eye-antennal disc development. For instance, Lola (Fig 2; clusters 2, 5, 7 and 12) regulates ocelli [53] and photoreceptor and cone cell development [54], Blimp-1 (Fig 2; cluster 2) controls the progression of retinal differentiation [55,56] and Mad (Fig 2; clusters 3, 4 and 15), the transcription factor that mediates Decapentaplegic (Dpp) – signalling [57], is involved in various processes in the eye-antennal disc [58–60]. Additionally, in 5 of 15 clusters (NES > 4) (Fig 2; e.g. clusters 7-9, 13 and 14) (in 9 of 15 clusters, NES > 3) we found a strong enrichment of motifs predicted to be bound by the GATA transcription factor Pnr, which is involved in establishing the early dorsal ventral axis of the eye [61–63]. Therefore, we revealed putative hub genes with important roles during the eye-antennal disc development, which regulate DEGs between *D. melanogaster* and *D. maurtiana.* Interestingly, 61.3 % (92 of 150) of the putative upstream transcription factors (i.e. putative hub genes) (NES > 4.0) were differentially expressed in at least one developmental stage (False discovery rate (FDR) 0.05) (Fig 2 and S2 Table). In summary, we identified many putative hub genes whose differential expression affected the transcriptomic landscape underlying eye-antennal disc development.

### Pannier is a phenotypically relevant hub gene in the GRN underlying dorsal head development

The observation that many of the putative hub genes were differentially expressed between species suggests that these changes may be causal in defining the observed differences in eye size and head shape. However, developmental processes [64,65] tend to be robust against perturbations because variation in the expression of developmental genes often does not result in phenotypic differences due to extensive buffering within GRNs [66–68]. Therefore, it remains questionable, whether the differential expression of the putative hub genes and thus the variation in expression of their target genes are indeed relevant for the differences in head morphology between *D. melanogaster* and *D. mauritiana*. To test this, we further investigated the role of Pannier (Pnr), which is an interesting putative hub gene for the following reasons: 1. Our global clustering and motif enrichment analyses suggest that Pnr regulates many DEGs between both species (Fig 2). 2. *pnr* itself is differentially expressed between *D. melanogaster* and *D. mauritiana*, with higher expression in the latter species (Fig 3A). 3. Pnr is known to be expressed in the dorsal eye-antennal disc [61] and it is involved in defining the dorsal-ventral axis of the eye [61,63,69]. Later during eye-antennal disc development, Pnr influences the ratio of retinal and head cuticle fate in the dorsal disc by repressing retinal determination genes [61,70,71].

**Fig 3.**
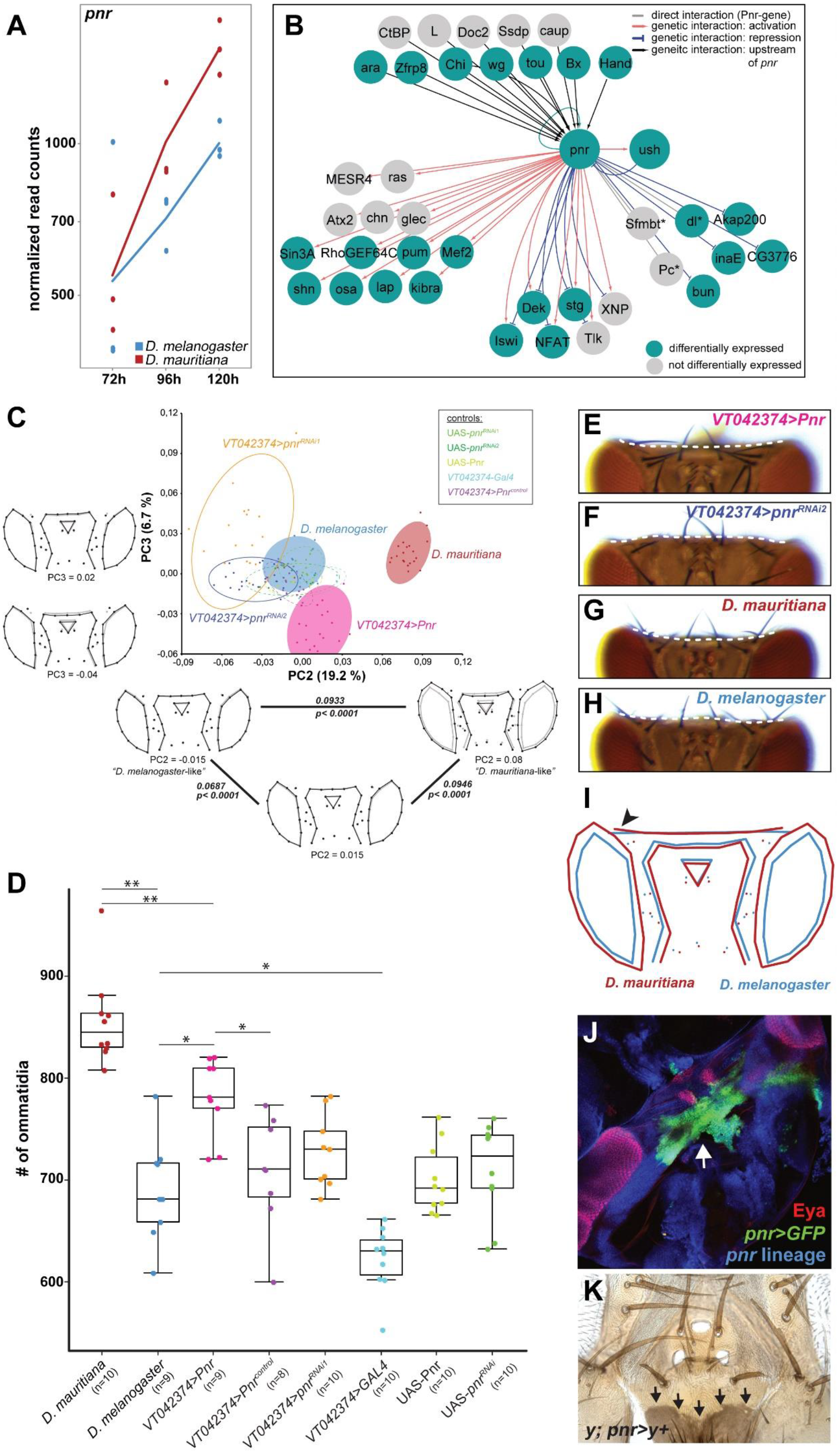
Pnr is a phenotypically relevant hub gene during eye-antennal disc development. **(A)** Expression dynamics of the *pnr* transcript at the three developmental stages in *D. melanogaster* (blue) and *D. mauritiana* (red) based on rlog transformed read counts. **(B)** Network reconstruction of known interactions upstream and downstream of Pnr that overlap with our Pnr target gene list. 14 of the 29 target genes are activated (red edges) and 8 genes are repressed (blue edges) by Pnr. 6 target genes showed both types of interactions. 21 of the 30 putative Pnr target genes (68%) were differentially expressed (green nodes). **(C)** Principal component analysis of dorsal head shapes. Shape outlines for PC values are given along the PC2 and PC3 axes, respectively, as indicated below each outline. In the triangle along PC2, the Procrustes distance between each shown shape outline is provided, as well as the p-value after permutation test with 1,000 cycles. Overexpression of *pnr* shifts the dorsal head shape towards a *D. mauritiana* head shape. **(D)** Number of ommatidia counted in one compound eye per individual. The significance was calculated by ANOVA and each pairwise comparison was calculated using Tukey’s HSD test; **<=0.01; *<=0.05. The color code in **(C)** and **(D)** is the same. **(E)** Representative dorsal view of an adult head after overexpression of *pnr* and **(F)** after knockdown of *pnr*. **(G)** Representative dorsal view of an adult head *D. mauritiana*. **(H)**Representative dorsal view of an adult head of *D. melanogaster*. The dotted lines in **E-H** mark the border of the occipital region. **(I)** Mean shapes of *D. melanogaster* (blue) and *D. mauritiana* (red) heads after discriminant function analysis including seven additional landmarks in the occipital region. The black arrowhead marks the lateral bending of the occipital region in *D. mauritiana*. **(J)** *pnr*-expression in developing pupal head structures. Cells marked with *pnr*>GFP accumulated in the future occipital region (green) behind the developing ocelli (red), and the head region where the two discs fuse (white arrow). **(K)** *pnr* lineage in adult dorsal heads shown by cells in which *yellow* expression is restored (black arrows).

To confirm that *pnr* is indeed a hub gene with many connections within the GRN orchestrating eye-antennal disc development, we first refined the list of its potential target genes. We searched for the Pnr-specific GATA motif in genomic regions that were accessible during eye-antennal imaginal disc development (see Materials and Methods and S1 Fig for details). In 14,511 open chromatin regions (across all timepoints) we found 1,335 Pnr-specific GATA motifs associated with 1,060 genes expressed in our RNAseq dataset, suggesting that those genes were active during eye and head development. In the regulatory regions of these 1,060 Pnr target genes we found a strong enrichment of Pnr binding sites (S4A Fig) and many of the genes were also predicted to be regulated by Pnr in our cluster analysis (see Fig 2 and S4B Fig). In line with the known functions of Pnr during eye-antennal disc development, the Pnr target genes were highly enriched in processes like compound eye development and growth as well as cell cycle progression (S4C Fig). Additionally, a cross-validation of our Pnr target genes with the DroID interaction database [72,73] showed that our list contained three known direct target genes (i.e. Pnr-regulatory sequence interaction; *dl, Pc* and *Sfmbt*) and 23 genes with known genetic interactions (Fig 3B), suggesting that these genes are also direct targets of Pnr. 17 out of these 26 genes (65.38 %) (Fig 3B) and 716 of the full set of 1,060 target genes (67.5 %) showed expression differences between *D. melanogaster* and *D. mauritiana* (S3 Table). Taken together, we confirmed that Pnr regulates many DEGs during eye-antennal disc development, suggesting that it is a hub gene with phenotypic relevance.

To test if variation in *pnr* expression can explain naturally occurring differences in eye size and dorsal head shape (Fig 1) we overexpressed and knocked down *pnr* in *D. melanogaster* in the dorsal eye-antennal disc in a domain that is reminiscent of endogenous *pnr* expression (S5A and 5B Fig). Successful manipulation of the expression was confirmed by reduced and enhanced protein expression in knock-down and overexpression discs, respectively (S5C-S5E Fig). Subsequently, we quantitatively compared the adult head morphology applying geometric morphometrics and ommatidia counting (see Materials and Methods for details). A principal component analysis of head shape variation showed that principal component 2 (PC2) explained 19.2 % of the observed variation in head shape and PC3 explained 6.7 % (Fig 3C; note that PC1 captured a technical artefact, see S6 Fig). Variation along PC2 mainly captured differences in the proportion of eye *vs.* interstitial cuticle in the dorsal head, as well as the location of the ocellar region. Intriguingly, the overexpression of *pnr* in the dorsal head region resulted in a shift from a “*D. melanogaster*”-like shape towards a more “*D. mauritiana*”-like shape along PC2, with an enlargement of the eyes at the expense of a slight reduction of the interstitial cuticle (Fig 3C). Ommatidia counting in entire eyes confirmed that the increase in eye area upon *pnr* overexpression was indeed due to an increase in number of ommatidia (Fig 3D).

PC3 explained mostly differences in the dorsal-posterior head cuticle and the location of the ocellar region (Fig 3C). We analysed the dorsal-posterior head cuticle in more detail and observed that the occipital region was more convex upon *pnr* overexpression (Fig 3E), whereas downregulation consistently led to an enlargement of these regions (Fig 3F). Intriguingly, in accordance with a higher expression of *pnr* in *D. mauritiana,* we found a more convex occipital region in *D. mauritiana* (Fig 3G) compared to a more concave shape in *D. melanogaster* (Fig 3H and 3I). Lineage tracing experiments showed consistent expression of *pnr* during eye-antennal disc development (S7 Fig) and detection of *pnr* expression in pupae (Fig 3J) as well as the analysis of *pnr*-expressing clones in adult heads (Fig 3K) confirmed that *pnr* was indeed expressed in the future occipital region. In summary, we showed that Pnr is a phenotypically relevant hub gene during eye-antennal disc development because we were able to phenocopy aspects of the “*D. mauritiana*”-like head shape and eye size by upregulating *pnr* expression in the developing eye-antennal disc of *D. melanogaster*.

### U-shaped modulates Pnr function during eye-antennal disc development

The detailed analysis of the heads of *pnr* gain- and loss of function flies revealed a consistent gain- and loss of vertical bristles, respectively (Fig 4A-4C). A similar role in sensory bristle development has been shown in the wing imaginal disc, where Pnr controls the positioning sensory bristles in the thorax [74–76]. The role of Pnr in thoracic sensory bristle development is antagonized by the presence of its co-factor U-shaped (Ush) [74,77]. Upon heterodimerization with Ush, Pnr that normally acts as a transcriptional activator, acquires a repressive function [77–79]. In line with a dual role of Pnr as transcriptional activator and repressor, we found genes that were activated as well as repressed by Pnr among its differentially expressed high confidence target genes during eye-antennal disc development (S8 Fig and Fig 3B). Additionally, the knock-down and overexpression of *ush* during eye-antennal disc development resulted in loss or misplacement of vertical bristles (Fig 4D and 4E), suggesting that Pnr and Ush may coordinate similar processes. Therefore, we asked whether Ush may also modulate the function of Pnr during eye-antennal disc development. To answer this question, we tested whether 1) Pnr and Ush are co-localized in the same cells in the eye-antennal disc and 2) both genes interact genetically.

**Fig 4.**
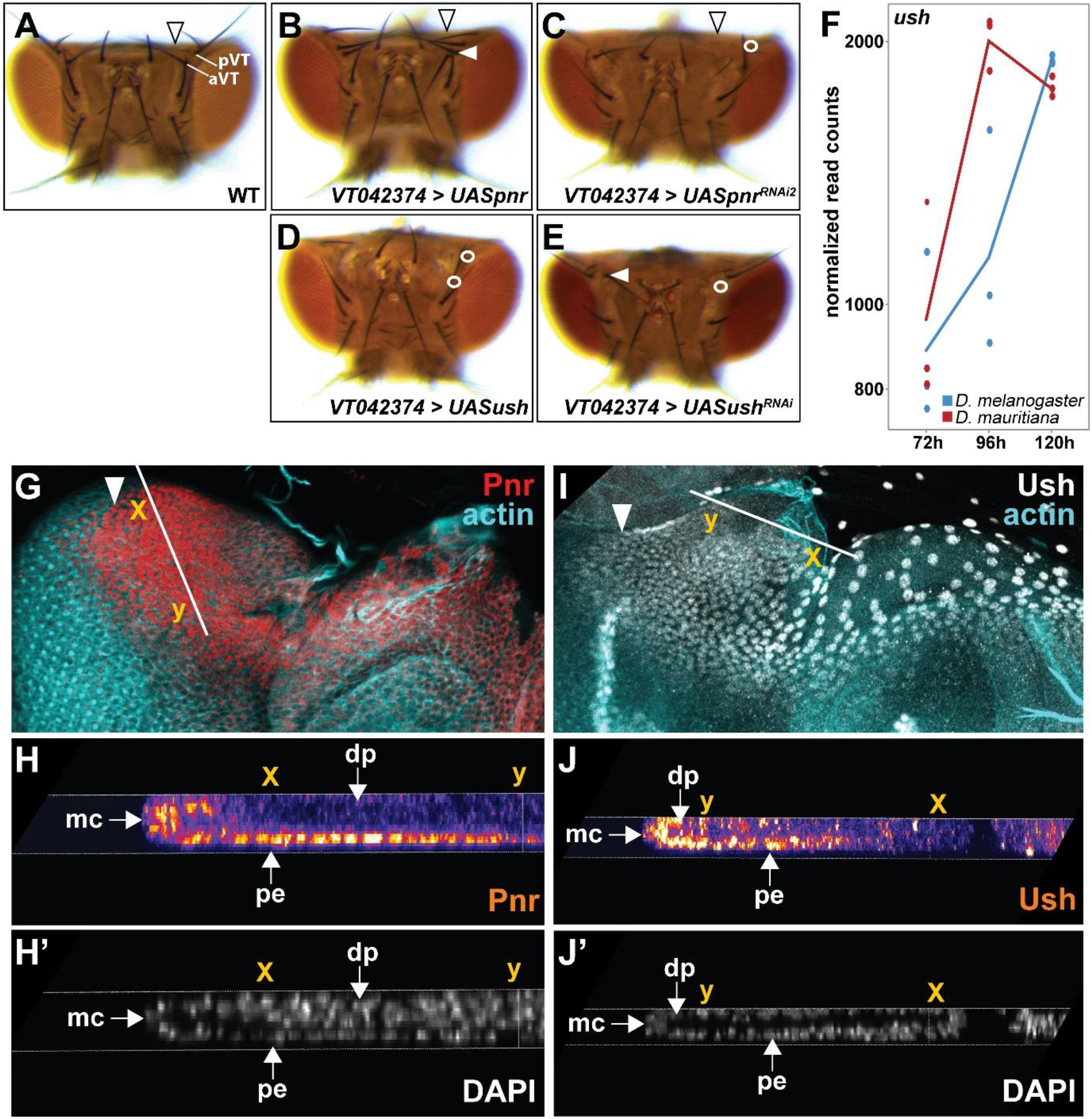
Pnr and Ush interact during eye-antennal disc development. **(A)** Dorsal view of an adult wild type head of *D. melanogaster*. Posterior (pVT) and anterior vertex bristles (aVT) are labelled. The arrowhead marks the occipital region. **(B)** Overexpression of *pnr* did not lead to major irregularities in the dorsal head cuticle, but to duplication of the pVT bristle next to the compound eye (white arrowhead) (N= 9 females, N=10 males). **(C)** Knock-down of *pnr* resulted in visible enlargement of the occipital region (black arrowhead) and to the loss of the pVT bristle next to the compound eye (white circle) (n= 10 females, 10 males). **(D)** Overexpression of *ush* resulted in irregularities in the dorsal head cuticle, and to the loss of the pVT and aVT bristles (white circles) next to the compound eye (n= 10 females). **(E)** Knock-down of *ush* resulted in slight irregularities of the dorsal head cuticle, to loss (white circle) or misplacement (white arrowhead) of the aVT bristle (n= 11 females). **(F)** Expression dynamics of the *ush* transcript at the three developmental stages in *D. melanogaster* (blue) and *D. mauritiana* (red) based on rlog transformed read counts. **(G, H)** Pnr protein location in 3rd instar eye-antennal discs in *D. melanogaster*. The Pnr protein is present in the dorsal peripodial epithelium (pe) of the developing disc (**G**), including a few cells of the margin cells (mc) and the disc proper (dp) (**H’’**). The white arrow in **G** marks the morphogenetic furrow, the solid white line marks region of the cross section shown in **H** and **H’** and the x and y coordinates indicate the same location in **G, H** and **H’**. **(I, J)** Ush protein location in 3rd instar eye-antennal discs in *D. melanogaster*. The Ush protein is, similarly to Pnr (compare to **G**), expressed in the dorsal peripodial epithelium (pe) of the developing disc (**I**), including a few cells of the margin cells (mc) and the disc proper (dp) (**J’**). The white arrow in **I** marks the morphogenetic furrow, the solid white line marks region of the cross section shown in **J** and **J’** and the x and y coordinates indicate the same location in **I, J**and **J’**.

Although, it was previously stated that *ush* is not expressed in the eye-antennal disc [61,78], we observed expression in all three studied stages in our RNAseq data (Fig 4F). As shown for *pnr* (Fig 3A), *ush* expression was mostly higher in *D. mauritiana* (Fig 4F). We confirmed the transcriptomics data using newly generated antibodies to show that both proteins are expressed in largely overlapping domains (Fig 4G-4J and S9 Fig) in the large nuclei of the dorsal peripodial epithelium (compare Fig 4G to 4I) and in a subset of the cuboidal margin cells in the disc proper (compare Fig 4H to 4J).

Since both proteins are co-localized in the eye-antennal disc, we tested for genetic interactions. We asked whether manipulation of *pnr* expression affected *ush* expression and vice versa. To this end, we overexpressed and knocked down both genes in the dorsal eye-antennal disc in a domain that is reminiscent of *pnr* expression (S5A and S5B Fig). We confirmed the successful overexpression and knock-down of both genes by enhanced and loss of protein expression, respectively (S5C-S5H Fig). Due to the qualitative nature of our experimental setup we could not obtain conclusive results for some of the tested combinations. However, we observed a reduced Ush signal upon knock-down of *pnr* (S5I Fig), suggesting a positive interaction between *pnr* and *ush*. Additionally, upregulation of *ush* resulted in slightly reduced Pnr protein levels (S5J Fig), suggesting a negative impact of Ush on *pnr* expression. Further support for this idea comes from the induction of a double-antenna phenotype when *ush* was overexpressed using a stronger *pnr* driver line (S5K Fig) [78], a phenotype that we also observed upon *pnr* knock-down (S5L Fig) [61]. In summary, we could show that Pnr and its co-factor Ush are co-expressed, interact genetically (S5M Fig) and are involved in dorsal sensory head bristle development. We suggest that a dual role of Pnr during eye-antennal disc development is most likely mediated by its co-factor Ush.

## Discussion

The trade-off between compound eyes and the interstitial cuticle in *Drosophila* heads is an excellent model to study the developmental and molecular mechanisms underlying complex trait evolution [22–24,39,42,44]. We established that this trade-off is present in the dorsal head of *D. melanogaster* and *D. mauritiana*, with the latter having larger compound eyes with significantly more ommatidia [this work and 42]. The comparison of larval eye-antennal disc development in both species revealed differences in the size of future head regions (i.e. retinal and antennal regions) starting at 96 h AEL, suggesting that variation in processes during late larval development are responsible for the observed differences in head morphology. To identify candidate genes driving these developmental differences, we combined comparative transcriptomics with chromatin accessibility data obtained prior to and after interspecific differences were observable (i.e. 72 h, 96 h and 120 h AEL). We show that the GATA transcription factor Pannier (Pnr) is a prime candidate hub gene: It showed higher expression in *D. mauritiana* and up to 67 % of its more than 1,000 predicted direct target genes were differentially expressed between species. In the following we propose a developmental model explaining how variation in *pnr* expression can simultaneously affect ommatidia number and interstitial cuticle size and we discuss our findings with respect to evolutionary hotspots and GRN evolution.

### Developmental mechanism underlying eye size and head shape differences between *D. melanogaster* and *D. mauritiana*

The spatially restricted overexpression of *pnr* in its endogenous domain resulted in a significant increase in ommatidia number, implying a direct effect on retinal development and thus final eye size. Our lineage tracing experiment showed that *pnr* positive cells contribute to the entire dorsal peripodial epithelium and to cells of the dorsal posterior margin where the morphogenetic furrow, and thus retinal differentiation, is initiated at late second/early third instar stages [80]. It has recently been shown that Eyeless/Pax6 (Ey) activity in the peripodial epithelium and in these posterior margin cells is necessary for morphogenetic furrow initiation and placement of the dorsal-ventral boundary [81]. The latter is tightly linked to reported functions of Pnr and tissue growth in the retinal region of the eye-antennal disc [61,63,69]. We found *ey* among the putative direct Pnr target genes, suggesting that Pnr may activate *ey* expression in the peripodial epithelium and in posterior margin cells. Since *ey* was not differentially expressed in our dataset, it remains to be tested if Pnr may regulate the final number of ommatidia clusters and thus final eye size through Ey-mediated initiation of differentiation and regulation of overall retinal growth prior to the stages studied here (Fig 5A).

**Fig 5.**
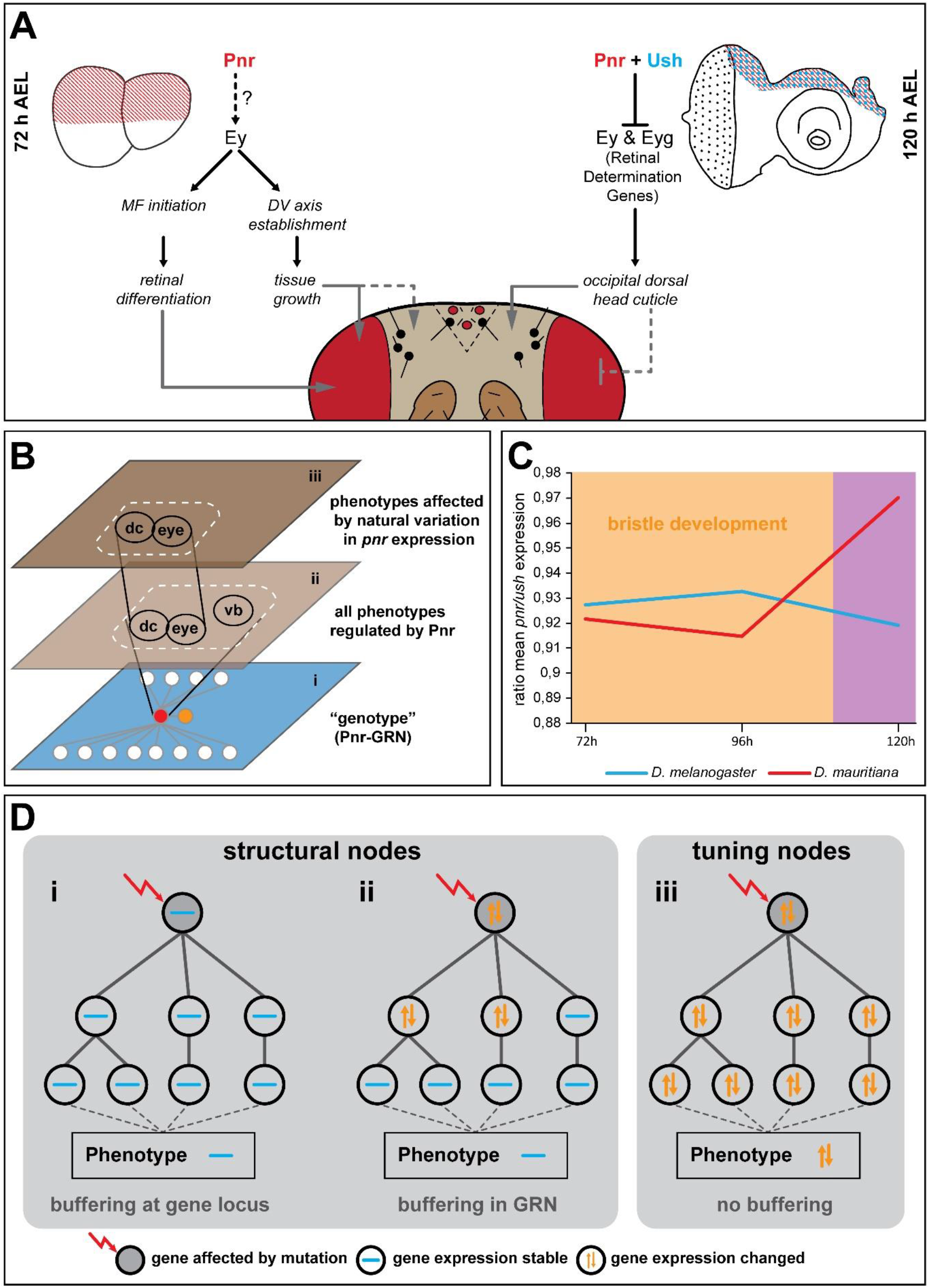
Developmental model and hypothesis about GRN evolution. **(A)** Summary of expression and functional data obtained in this work. Black connections indicate genetic interactions among genes. Grey connections represent observed phenotypic effects upon manipulation of gene expression. **(B)** Schematic representation of functional constraints on vertical bristle development. The genotype level (i) is depicted as GRN centred on Pnr (red node in i). Pnr regulates the development of the dorsal occipital cuticle (dc), the compound eye (eye) and vertical bristles (vb) (ii). Due to a functional constraint on vertical bristle development only the effect of differences of *pnr* expression on dc and eye are phenotypically relevant during the evolution of head morphology (iii). We propose that the constraint effect of *pnr* expression differences between *D. melanogaster* and *D. mauritiana* is mediated by co-evolution of *ush* expression (orange node in i). **(C)** The ratio of *pnr* and *ush* expression is stable at 72 h and 96 h AEL when vertical bristles are defined. Later expression of both genes is not constraint anymore by a function in bristle development. **(D)** Schematic representation of structural (i, ii) and tuning nodes (iii) in developmental GRNs.

We found that *pnr*-expressing cells contribute to the dorsal interstitial head region in pupae, suggesting that the phenotypic effects in the occipital head region most likely highlight late functions of Pnr. Indeed, Pnr represses members of the retinal determination network throughout the third instar stage to regulate the ratio of retinal cells *vs*. head cuticle cells [71]. In addition to *ey,* we found *eyegone* (*eyg*) as second member of the retinal determination network among putative direct Pnr targets. Both genes must be repressed to ensure proper dorsal interstitial cuticle development [82,83], suggesting that Pnr may act through transcriptional repression during late dorsal head development. Such a repressive role could be mediated upon heterodimerization with its co-factor Ush as shown for the wing disc [70,77,84,85]. Indeed we observed that, despite previous reports [61], the Ush protein is co-localized with Pnr in dorsal peripodial epithelium and in the cuboidal cells of the disc margin. In accordance with their co-expression, both genes interact genetically and they are both necessary for proper dorsal head development and the formation of the same head bristles (this work and see [74] for bristle phenotype). Therefore, we propose that Ush mediates a repressive late function of Pnr in the dorsal peripodial epithelium to regulate retinal *vs.* dorsal head cuticle development during the third larval instar stage (Fig 5A).

Taken together, our data shows that variation in *pnr* expression during late larval development can simultaneously affect the size of the dorsal interstitial head cuticle and the compound eyes (Fig 5A). Accordingly, increasing *pnr* expression levels in *D. melanogaster* mimics higher *pnr* expression as observed in *D. mauritiana* and phenocopied major aspects of the dorsal *D. mauritiana* head shape and eye size. Consistent with the fact that variation in complex traits is caused by multiple loci [15,17], *pnr* overexpression was not sufficient to fully phenocopy *D. mauritiana* head shape and eye size. Furthermore, it remains to be established whether the *pnr* and/or *ush* loci contain genetic variants associated with eye size and head shape differences. While quantitative genetics approaches are not applicable due to the infertility of F1 females in interspecific crosses between *D. melanogaster* and *D. mauritiana* [86], reciprocal hemizygosity tests [87] for Pnr, Ush and putative regulators of these two factors represent a powerful tool to further dissect the causative genetic variants in the future.

In summary, based on our developmental model the increase in eye size in *D. mauritiana* that goes along with a reduction of dorsal interstitial cuticle and a convex bending of the occipital head region can partially be explained by natural variation in *pnr* expression. Since Pnr is a highly connected pleiotropic transcription factor that regulates many differentially expressed genes between *D. melanogaster* and *D. mauritiana*, we argue that Pnr is a phenotypically relevant hub gene underlying natural variation in head morphology.

### Functional constraints drive co-evolution of *pnr* and *ush* expression

In line with the pleiotropic role of Pnr during eye-antennal disc development, classical gain- and loss of function experiments resulted in extensive phenotypic consequences ranging from retinal overgrowth to the entire loss of the compound eyes [61,71]. These rather coarse experiments based on highly artificial systems, such as ectopic expression of *pnr* induced by an *eyeless/Pax6*-Gal4 driver line that is active from early stages on in most of the eye-antennal disc [81], are suitable to unravel the combined functions of a pleiotropic gene. However, it is a major challenge to disentangle the distinct tissue and stage specific functions of pleiotropic factors and find out which of these are phenotypically relevant in an evolutionary context [35,88]. This is particularly relevant because developmental processes [64,65] and the expression of the genes regulating them [89–91] tend to be robust against genetic or environmental perturbations. Therefore, functional and developmental constraints limit the evolutionarily relevant phenotypic effects of pleiotropic genes (Fig 5B).

We observed an excellent example for such a constraint. While individual gain- and loss of function of *pnr* and *ush* in *D. melanogaster* affected the formation of anterior and posterior vertical sensory bristles in the dorsal head, we never observed gain or loss of these bristles between species. Pnr and Ush have been shown to be involved in thoracic sensory bristle development by controlling the positioning of proneural cell clusters in the wing disc [74–76]. Thoracic sensory bristles develop in regions with high *pnr* and low *ush* expression [74,77] and their balanced expression is facilitated by genetic interactions between both genes [84]. We observed that *pnr* overexpression in the eye-antennal disc consistently resulted in duplication of the anterior vertical bristles, suggesting that extra bristles arise where *pnr* overexpression cannot be compensated by Ush [see also 76]. Similarly, overexpression of *ush* in most of the dorsal peripodial epithelium resulted in loss of the anterior and posterior vertical bristles, showing that extra Ush above a certain threshold completely antagonizes Pnr function and subsequent sensory bristle formation. Hence, the stoichiometry between Pnr and its co-factor Ush is crucial for proper sensory bristle development in the dorsal head as well.

How can natural variation in *pnr* and *ush* expression cause differences in head morphology without affecting the sensory bristle pattern? Since most of the proneural cell clusters are specified 18h to 6h prior to puparium formation [92], the ratio between Pnr and Ush must remain stable at 72 h and 96 h AEL to ensure proper sensory bristle formation. However, once bristles are specified, variation in the ratio of both factors at later stages (e.g. 120 h AEL) can cause natural variation in the shape of the dorsal occipital head region without affecting bristle formation (Fig 5C). Accordingly, both genes showed higher expression in *D. mauritiana* at 72 h and 96 h AEL resulting in a similar ratio of both factors (Fig 5C). In contrast, a different ratio was observed at 120 h AEL, with *pnr* expression being higher in *D. mauritiana*, while *ush* expression was more similar in both species at this later stage (Fig 5C). We propose that the functional constraint on stereotypic sensory bristle development drove the co-evolution of *pnr* and *ush* expression during early third instar disc development in *D. melanogaster* and *D. mauritiana* (Fig 5C). Indeed, according to their crucial function in environmental perception [93], vertical bristles have been shown to be robust against temperature fluctuations during development [94]. Therefore, the context-dependent presence and action of a transcriptional co-factor provides a versatile mechanism to restrict the phenotypic effects of a pleiotropic developmental factor in an evolutionary context (Fig 5B).

### Repeated evolution of hotspot genes may play a minor role for complex trait evolution

Natural intra- and interspecific variation in eye size and a trade-off between retinal tissue and head cuticle is a common feature of *Drosophila* [22,24,39,41,42,44,48]. The most comprehensive genetic and developmental characterization of this trait suggested that differences in the early subdivision of the eye-antennal disc driven by variation in the temporal regulation of *eyeless/Pax6* expression is a common mechanism driving the trade-off [44]. In line with the identification of evolutionary hotspot genes for rather simple traits, such as variation in pigmentation patterns [10,11] and the gain and loss of structures [12–14], this finding suggests that repeated natural variation in hotspot genes may be a major driver for the evolution of complex morphological traits as well [44]. In contrast to these results, our comparison of eye-antennal disc development between *D. melanogaster* and *D. mauritiana* showed that differences in the anlagen of the future retinal field and the antenna arise only late during the third instar stages. Accordingly, our developmental model involving natural variation in *pnr* and *ush* expression (Fig 5A) provides an alternative developmental mechanism that can explain the simultaneous effects on ommatidia number variation and differences in the interstitial head cuticle.

Recent advances in studying the developmental and genetic basis of the trade-off between eye and interstitial cuticle size in *Drosophila* revealed that differences in developmental processes affecting both traits simultaneously seem to be common in *Drosophila* [23,24]. However, eye and interstitial cuticle size can also be genetically and developmentally uncoupled as shown by a detailed characterization of the head trade-off in *D. melanogaster* and *D. simulans* [22]. Quantitative genetics data for intraspecific variation in the head trade-off in *D. melanogaster* and *D. simulans* shows that its genetic basis is highly diverse [23], supporting our observation that repeated evolution through hotspot genes may not be prevalent in the evolution of head morphology.

The insect compound eye is composed of hundreds of functional subunits, the ommatidia, and they themselves are built by 14 unique cells [95]. Therefore, the size of an insect compound eye can vary due to the number of ommatidia or due to their size. Indeed, a detailed morphological analysis showed that differences in the number of ommatidia explains intra- and interspecific variation in various *Drosophila* species [this work and 23,42,44], while eye size differences between *D. simulans* and *D. mauritiana* are caused by variation in ommatidia size [22,42]. Since these two features are regulated at different time-points by different developmental processes, it is expected that the molecular and developmental basis of eye size differences varies in different groups. Therefore, depending on the cellular basis of observed morphological differences, different developmental and molecular mechanisms can affect complex trait evolution.

A potential explanation for the convergent evolution of the head trade-off may be the complex modular nature of head development. The eye-antennal disc comprises compartments for all future head structures, including the eye, the interstitial cuticle, and the antennae. Besides the already established mechanisms [this work and 44], it is conceivable that differences in adult head morphology can be the result of variation in different developmental processes in each compartment of the disc at different time points. Additionally, the development of a complex organ, such as the insect compound eye, is controlled by several hub genes. This is best exemplified by the observation that the experimental mis-expression of different members of the retinal determination network can induce an ectopic retinal developmental program [96– 100], suggesting that they all act as input-output devices in the retinal GRN. In contrast, the development of rather simple morphological traits is regulated by very few hub genes that act as input-output devices integrating positional information and activating most target genes required to initiate the entire developmental program to form the structure [33,34]. In summary, our current knowledge based on quantitative genetics, developmental as well as morphological data suggests that different nodes within the GRN underlying head and eye development evolve to give rise to variation in head morphology in *Drosophila*. We argue that the complexity and modularity of complex organ development facilitates the convergent evolution of complex morphological traits.

### Context dependent regulatory interactions facilitate the evolution of pleiotropic hub genes

Although we did not find indications for repeated evolution at hotspot genes for natural variation in eye size and head shape in *Drosophila*, our data and the work of others consistently revealed variation in the expression and function of crucial pleiotropic genes contributing to morphological evolution [e.g. this work and 6,7,44]. On the first glance this observation is counter-intuitive for two reasons: 1) The mutation rates of central developmental regulators, such as transcription factors and signalling molecules, are low [101–103] and such factors tend to be functionally conserved across the animal kingdom [97]. 2) Systematic analyses of GRN connectivity showed that developmental regulators are often highly connected [104,105]. Since GRNs are thought to be scale-free with few highly connected hub genes, mutations are more likely to occur in the many weakly connected peripheral genes [15,30]. Therefore, the question arises, why morphological evolution is often driven by natural variation in expression of highly connected pleiotropic genes?

First, although a high level of network connectivity ensures network stability, data obtained in yeast showed that genes that are regulated by many other factors (*trans*-mutational target size) and genes that contain many transcription factor binding motifs (*cis*-mutational target size) are more prone to accumulate mutations [90,106]. A study of Mendelian and complex disease genes revealed that they are often highly connected and that they act as brokers, connecting genes that are themselves poorly connected. Such broker positions could represent fragile nodes and variation in such genes may more likely result in phenotypic differences [107]. Although it remains to be tested if highly connected developmental regulators occupy similar fragile positions within GRNs, the position of pleiotropic genes within GRNs may influence the evolvability of their expression or function.

Second, highly connected genes often contain a complex regulatory landscape with many *cis*-regulatory elements allowing for their incorporation in different developmental GRNs. Consequently, a complex gene regulation facilitates the remodeling of regulatory interactions in a temporally and spatially defined manner [reviewed in 108]. Eye-antennal disc development is highly complex and the regulatory interactions within the underlying GRN are variable both throughout time [49] and in different parts of the disc [37], requiring the use of the same developmental gene products in different contexts. For instance, genes of the retinal determination network are essential for the initial proliferation and growth of the entire eye-antennal disc [56,109,110] and later they play a pivotal role in retinal specification [80,111]. These distinct roles have been suggested to be achieved by considerable rewiring of the respective GRNs, which allows them to fulfil temporally and even spatially restricted tasks [112]. The various described roles for Pnr during eye development [61,71], its continuous expression in the dorsal eye-antennal disc and the observation that variation in *pnr* expression affects overall head shape and eye size simultaneously (this work), strongly suggest that Pnr as well is involved in several sub-networks during eye and head development. The interaction with co-factors, such as Ush provides a mechanism of modulating the role of Pnr from an activating to a repressing transcription factor and its usage in spatially defined GRNs. Hence, the context-specific modulation of expression of a highly pleiotropic developmental factor, such as *pnr* and its target genes is facilitated by complex and modular gene regulation mechanisms and a highly dynamic underlying GRN.

### A versatile method to identify tuning nodes within developmental GRNs underlying morphological evolution

Since natural variation in gene expression is a major driver of morphological divergence [33,113], many studies aiming at revealing candidate genes underlying the evolution of morphological traits employ comparative transcriptomics methods to study differential gene expression. However, changes in gene function and expression are often buffered in GRNs to maintain a stable phenotypic outcome [68,114–119]. This is best exemplified by the observation that the individual loss of function of many genes in yeast [116] and the nematode *Caenorhabditis elegans* [118] rarely has deleterious effects for the organism. In accordance with a generally conserved adult head morphology in *D. melanogaster* and *D. mauritiana*, we found that the expression of many genes representing key developmental and metabolic processes was conserved between the two species. However, the observed quantitative interspecific differences in eye size and head shape must be due to variation in some parts of the GRN which are then selected for if it translates into phenotypic diversity that is advantageous for the organism [120–122].

In the light of the observed discrepancy between a generally high network stability and the flexibility in some parts of the network, we propose that developmental GRNs must contain at least two types of nodes (i.e. genes): structural nodes and tuning nodes (Fig 5D). Structural nodes confer phenotypic stability because variation in such nodes (e.g. variation in expression of the gene) is either prevented by tight regulation at the locus (Fig 5Di) or buffered downstream in the GRN (Fig 5Dii). Such a robustness has been demonstrated to be often the result of redundancies [114], either on the level of *cis*-regulatory elements [115,119] or transcription factors and signaling molecules [68,117]. In contrast, tuning nodes represent genes for which variation is tolerated within the GRN and results in quantitative phenotypic variation (Fig 5Diii). Natural variation in *ey* [44] and *pnr* expression (this work), which translates into head shape and eye size differences, as well as the previously identified hotspot genes underlying variation in trichome formation [12] and body pigmentation [10,11] are excellent examples of such tuning nodes. It is important to note here that developmental genes may represent tuning nodes in some contexts, while they are structural nodes in most other contexts. Our observation that interspecific differences in *pnr* expression affected the dorsal head shape and eye size, but not sensory bristle formation is an excellent example for the context-dependence of tuning and structural nodes. Revealing structural and tuning nodes in developmental GRNs allows gaining new insights into constrained and variable developmental processes, respectively.

Since many highly connected hub genes with pleiotropic functions have been shown to regulate morphological differences, they can be identified based on their effect on their target genes [36]. A systematic analysis of GRN properties suggests that transcription factor-to-gene interactions are scale-free, meaning that some transcription factors regulate a high number of genes [30]. Therefore, we hypothesized that putative tuning nodes (e.g. genes coding for transcription factors) can be elucidated by their impact on the expression of their target genes [i.e. “gene modules”; 50–52]. Accordingly, our methodological framework combines the identification of DEGs, clustering of DEGs based on their developmental expression dynamics and the identification of putative shared upstream regulators integrating genome wide chromatin accessibility data (Fig 2). Since fluctuations in gene expression are often buffered within GRNs, not all expression differences of putative tuning nodes will result in phenotypic differences. Therefore, a functional validation is crucial to reveal phenotypically relevant tuning nodes. Such a validation can be achieved by applying classical developmental genetics methods as shown in this work. If such tools are not established, a tuning node could be validated using a hemizygosity test based on widely applicable CRISPR/Cas9 aided genome editing [87]

Our approach unfolds its full potential if complemented with quantitative genetics data to identify exact genetic variants constituting the tuning node. This is relevant because mutations affecting the expression or function of tuning nodes could be in its locus itself or it could be the result of genetic changes in an upstream regulator. In the latter case it is expected that the number of potential candidates upstream of the tuning node is low because systematic analyses of GRN architecture revealed that the in-degree within networks follows a random distribution, suggesting that most transcription factors tend to be regulated by only a few upstream factors [30]. Therefore, once putative tuning nodes are identified by comparative transcriptomics, the number of candidate genes that regulate the tuning node is expected to be low facilitating further molecular characterization.

In summary, our comparative transcriptomics approach can be used as entry point to study the evolution of complex morphological traits or it can be applied to link already identified genetic variation to nodes within developmental GRNs and to developmental processes. Since the individual steps are applicable in many organisms, including those for which quantitative genetics approaches are not applicable, we are convinced that the approach represents a versatile framework to study the molecular and developmental basis of morphological evolution.

## Material and Methods

A detailed description of all methods is available as Extended Materials and Methods in the Supporting Information.

### Comparative Transcriptomics and ATACseq

Eye-antennal imaginal discs were dissected 72h, 96h and 120h after egg laying (AEL) from larvae of *D. melanogaster* (Oregon R) and *D. mauritiana* (TAM16), respectively. Total RNA was extracted from the discs and libraries for Illumina HiSeq2000 sequencing were generated using the TruSeq RNA Sample Preparation Kit (Illumina, catalog ID RS-122-2002). Reads were mapped using Bowtie2 v. 2.3.4.1 [123], mapping data was processed using samtools version 1.9 [124,125] and differential expression analysis was performed with the DESeq2 package (DESeq2_1.22.2 [126]; R version 3.5.2). Metascape was used to analyse differential enrichment of GO terms for pairwise comparisons. Clustering of genes according to their developmental expression dynamics was performed with the coseq package (version 1.6.1) [127,128] and we searched for potential upstream factors using the i-cisTarget tool [129,130]. ATACseq to assess chromatin accessibility was perfromed as described before [131]. The pipeline used to define a high confidence target gene list of Pnr is described in detail in the Extended Materials and Methods.

### Genetic crosses, Immunohistology and phenotyping

All fly crosses to knock-down and overexpress *pnr* and *ush*, as well as crosses for lineage tracing experiments were performed at 25°C and at a constant 12h:12h light:dark cycle. We generated polyclonal antibodies against Pnr [132] and Ush [78] based on previous knowledge (Proteintech, Rosemont, IL, USA) and immunohistology was performed as described in the Extended Materials and Methods. Adult heads were either mounted in Hoyer’s medium:Lactic Acid (50:50) or directly imaged for ommatidia counting or shape analysis using Geometric Morphometrics.

## Acknowledgments

Many thanks to Marita Büscher and Kolja Eckermann for sharing antibodies and to Alistair P. McGregor, Marc Haenlin and Gerd Vorbrüggen for sharing *Drosophila* stocks. We are grateful for financial support for the ATACseq data, as well as for critical comments on an earlier version of this manuscript to Alistair P. McGregor. Stocks obtained from the Bloomington Drosophila Stock Center (NIH P40OD018537) were used in this study. Many thanks to the Deep-Sequencing Core Facility of the Universitätsmedizin Göttingen (UMG) for next generation sequencing. We are grateful for many fruitful discussions and feedback from members of the Department of Developmental Biology and Posnien Lab members.

