## Supporting Information for "Variation in a pleiotropic hub gene drives morphological evolution: Insights from interspecific differences in head shape and eye size in *Drosophila*"

**This PDF file includes:**

S1 – S10 Figures

S1 Table. Number of differentially expressed genes (DEGs).

Extended Materials and Methods

**The following Supporting Tables are available separately:**

S2 Table. GO enrichment and iCis-Target analyses for clusters of differentially expressed genes.

S3 Table. List of putative Pnr target genes.

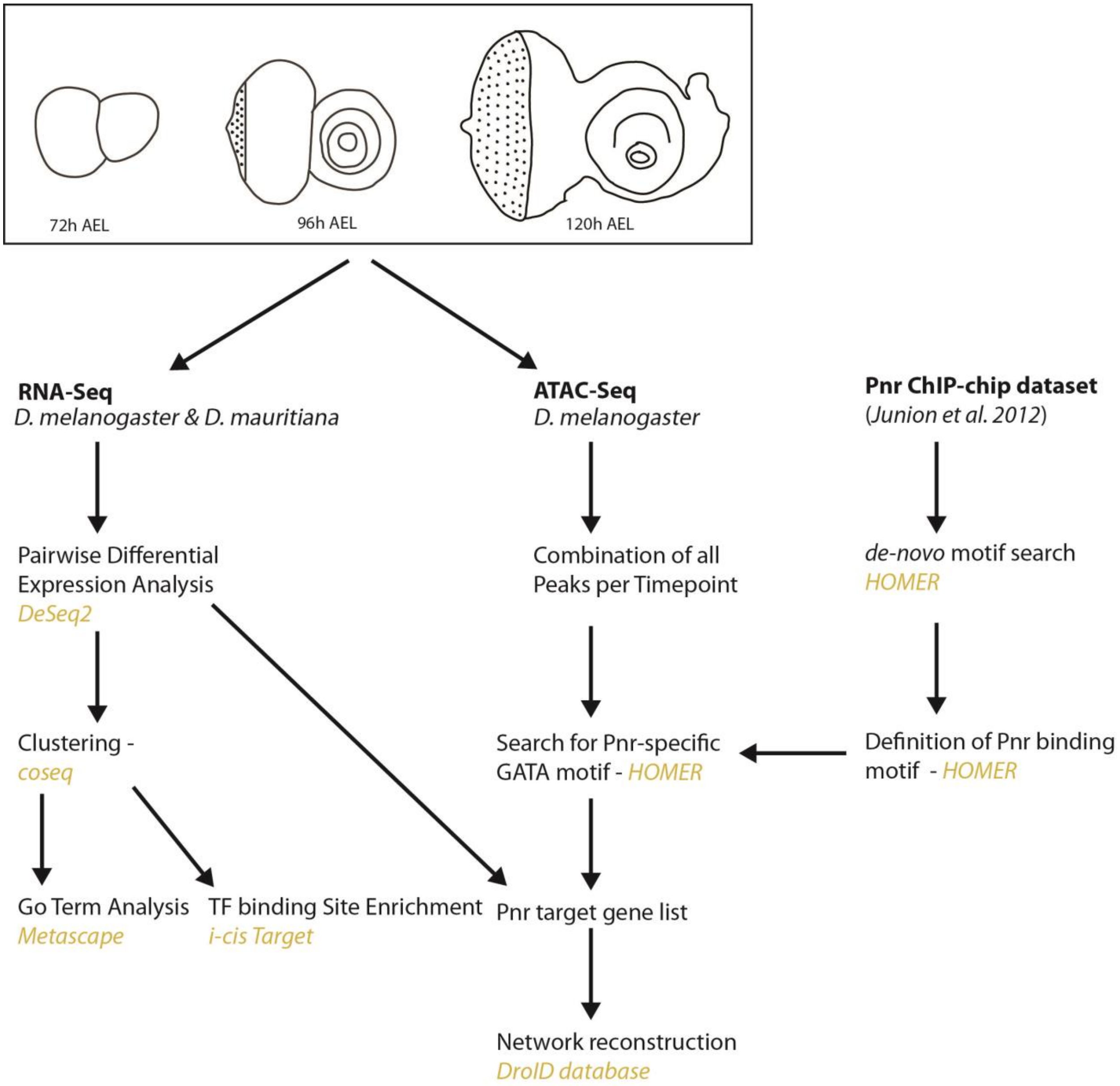


**S1 Fig. Overview of RNAseq and ATACseq analyses.** Scheme of the datasets generated and used for the analysis, together with the analysis pipeline. We generated RNA-seq and ATAC-seq datasets of developing eye-antennal discs at three stages at late-L2, mid-L3 and late L3 (72h AEL (after egg laying), 96h AEL and 120h AEL). The arrows outline the analysis pipeline and programs used in each step are given in yellow.

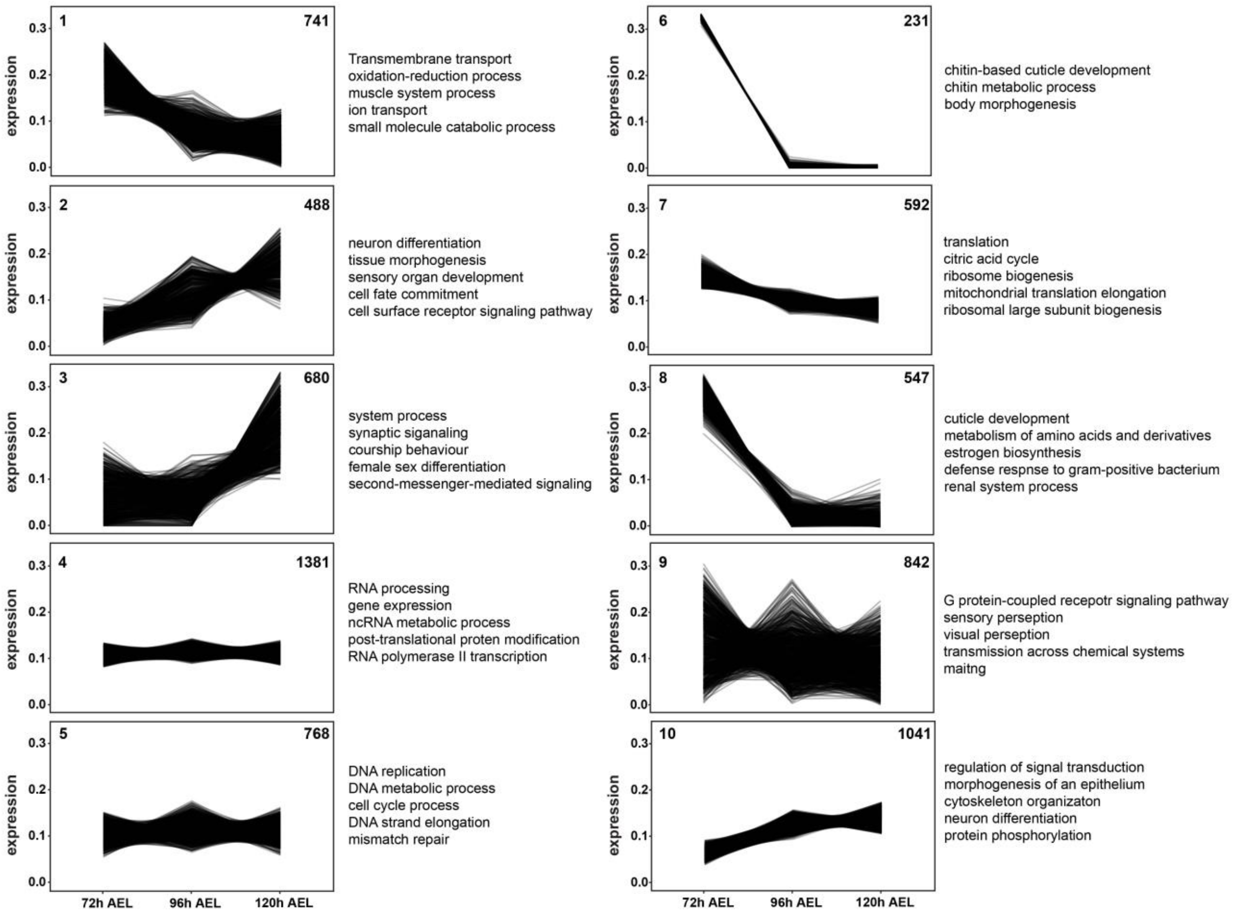


**S2 Fig. Clustering of genes that showed conserved expression between *D. melanogaster* and *D. mauritiana*.** Conserved genes were clustered based on expression dynamics across the three stages and a GO enrichment analysis was performed for each cluster. The number of genes in each cluster is given in the top right corner. Genes that were upregulated at early stages or constantly expressed over the three time-points, are mainly enriched in metabolic processes, cell-cycle processes and gene expression (e.g. clusters 4, 5 and 7). This is consistent with the fact, that the developing eye-antennal disc is a highly proliferative and growing tissue [1]. Genes that were highly expressed at 72h AEL and barely expressed at the two later stages, are enriched in cuticle development (e.g. clusters 6 and 8), reflecting the moulting of the *Drosophila* larva at the end of the 2$^{nd}$ instar. Genes that got steadily upregulated at the two later stages are mainly enriched in neuronal processes (e.g. clusters 2,3 and 10), consistent with the ongoing differentiation of ommatidia at these time-points [2].

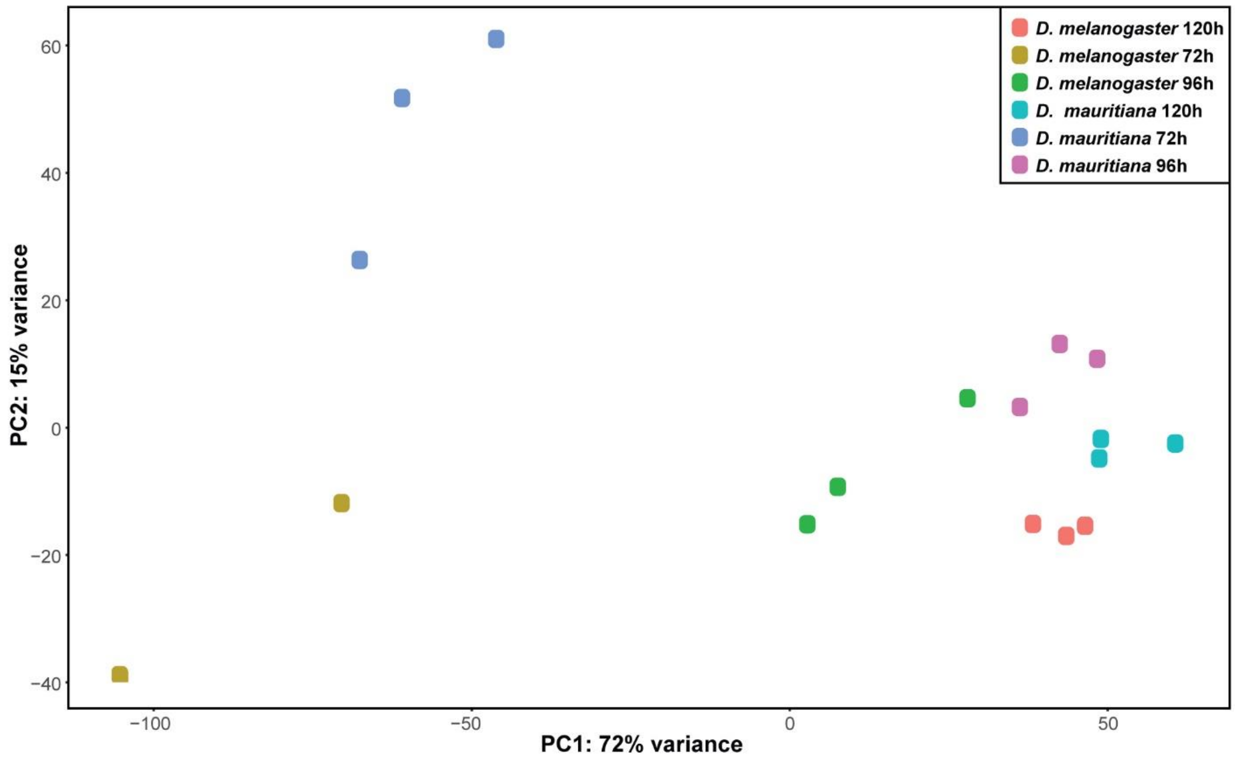


**S3 Fig. Principal Component Analyses of all RNAseq samples.** Analysis is based on rlog transformed read counts. PC1 separates the samples according to time-points, whereas PC2 separates them mainly the by species.

**A**

| possibleTFs | NES |
|---|---|
| pnr | 7.13 |
| nejire | 5.71 |
| Pmad | 4.75 |
| mef2 | 3.43 |

**B**

| Cluster_Number | ClusterSize | pnr_enriched?_Rank | NES | OverlapHCL | OverlapHCL% |
|---|---|---|---|---|---|
| 1 | 272 | 137 | 3.1 | 3 | 1% |
| 2 | 458 | 94 | 3.4 | 8 | 2% |
| 3 | 354 | 61 | 3.5 | 5 | 1% |
| 4 | 488 | - | - | 42 | 9% |
| 5 | 224 | - | - | 1 | 0% |
| 6 | 508 | - | - | 10 | 2% |
| 7 | 582 | 33 | 4.4 | 15 | 3% |
| 8 | 804 | 11 | 6.9 | 76 | 9% |
| 9 | 488 | 31 | 9.5 | 60 | 12% |
| 10 | 346 | - | - | 10 | 3% |
| 11 | 654 | - | - | 89 | 14% |
| 12 | 392 | - | - | 10 | 3% |
| 13 | 912 | 70 | 5.2 | 215 | 24% |
| 14 | 1499 | 3 | 18.9 | 168 | 11% |
| 15 | 369 | 209 | 3.1 | 4 | 1% |

**C**

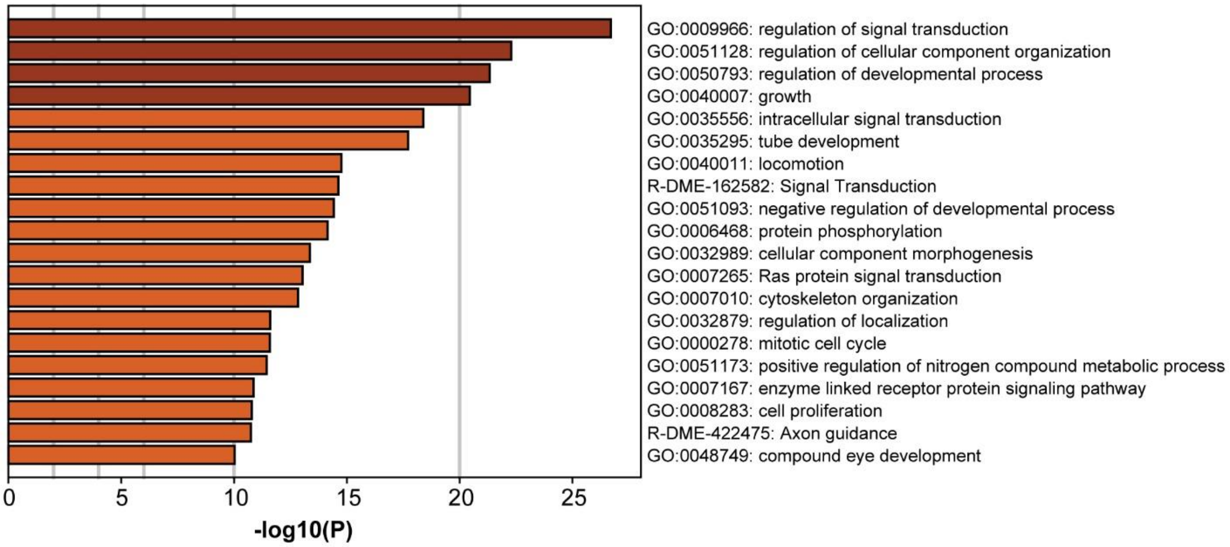


**S4 Fig. Cross validation of putative direct Pnr target genes. (A)** Transcription factor binding site enrichment in 5kb upstream of the transcription start site, 5’UTR regions and 1st introns of all predicted Pnr target genes (NES-values represent enrichment score). The identification of putative pMad target genes among Pnr targets may confirm the previous observation that both proteins interact physically during larval development [3]. **(B)** Table showing the number of DEGs clustered together, and if available, enrichment of the Pnr motif based on i-cis Target, together with the NES score. The last two columns show the overlap between the predicted Pnr target gene list and the respective cluster. **(C)** GO-term enrichment analysis of all predicted Pnr target genes. Enrichment in processes such as signal transduction, growth, cell cycle but also in more specific terms like compound eye development were identified.

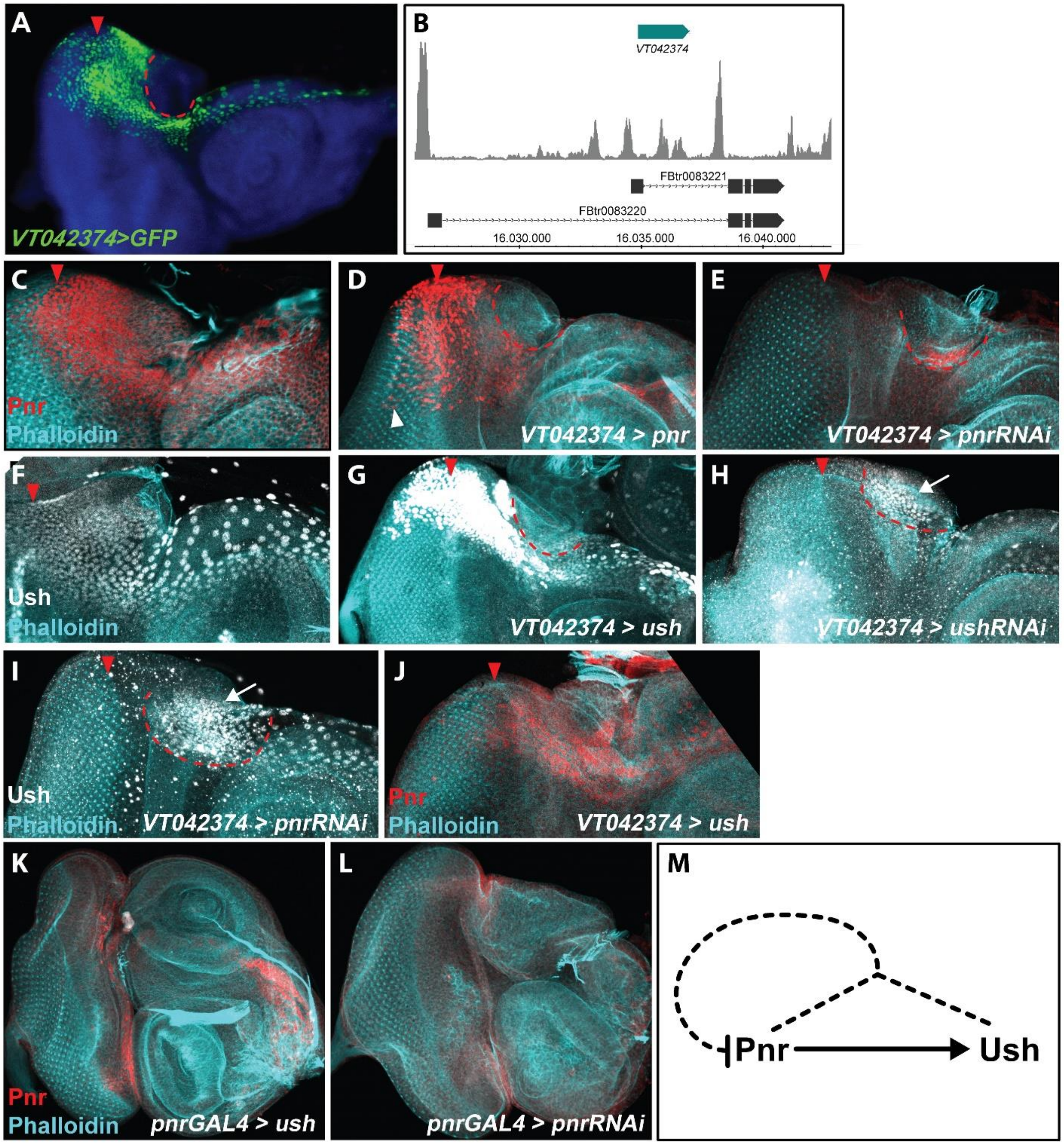


**S5 Fig. Genetic interactions of *pnr* and *ush* in *D. melanogaster*. (A)** The VT042374 lines drives expression of GFP in the dorsal peripodial epithelium. Note that the line does not drive expression in the ocelli region (red dotted line), where endogenous Pnr expression is weaker (compare to **c**). Furthermore, the GFP expression extends more posteriorly beyond the morphogenetic furrow (red arrowhead) compared to the wildtype protein expression of Pnr (compare to **c**). **(B)** Analysis of the *pnr* locus showed that the enhancer fragment of the VT042374 line overlaps with open chromatin (ATACseq peaks; grey) regions in the first intron, strongly suggesting that partial endogenous *pnr* expression is reported. **(C)** Wildtype Pnr expression detected with anti-Pnr antibody. **(D)** Overexpression of *pnr* with the VT042374 line leads to ectopic expression of Pnr posterior to the morphogenetic furrow (white arrowhead). **(E)** *pnr* RNAi knockdown with the VT042374 line resulted in strong reduction of Pnr expression. **(F)** Wildtype expression of Ush detected with anti-Ush antibody. **(G)** Overexpression of *ush* with the VT042374 line leads to increased expression of Ush as well as ectopic expression posterior to the morphogenetic furrow. **(H)** ush RNAi knockdown with the VT042374 line resulted in strong reduction of Ush expression in most of the dorsal peripodial membrane except for the ocellar region, where the VT042374 line does not drive expression (white arrow). **(I)** Knockdown of *pnr* resulted in reduction of Ush expression, except for the ocellar region, where the VT042374 line does not drive expression (white arrow). **(J)** Overexpression of *ush* resulted in reduction of Pnr expression. **(K)** Overexpression of *ush* resulted in duplication of antennae anlagen. **(L)** Knockdown of *pnr* resulted in duplication of antennae anlagen. **(M)** Summary of putative regulatory interactions between *pnr* and *ush*.

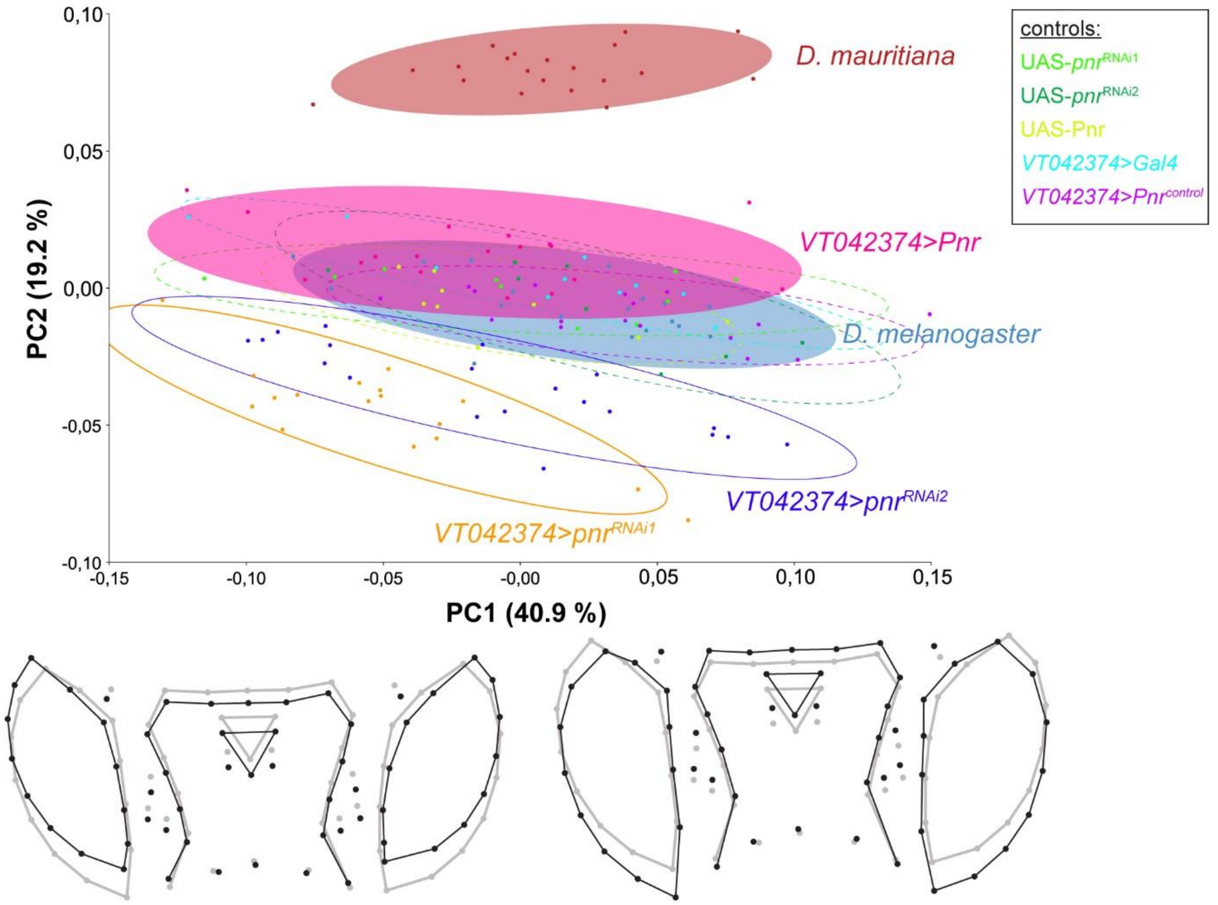


**S6 Fig. Principal Component Analysis showing artificial shape changes due to variation in positioning of the heads during image acquisition.** PC1 captured up- and down facing heads (see shape outlines representing -0.1 and 0.1 along PC1). Since this effect is present across all samples, this positioning artefact affects all samples simultaneously. We excluded the first principal component (PC1) and analyzed PC2 and PC3 in more detail.

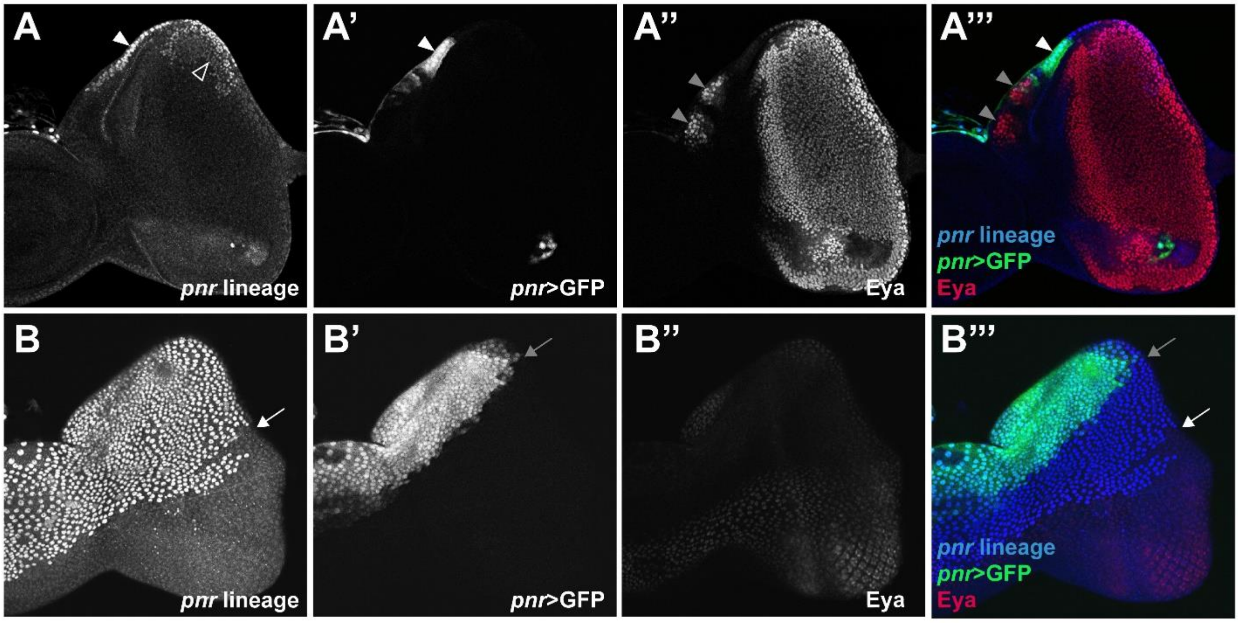


**S7 Fig. Lineage of *pnr*-expressing cells in the developing eye-antennal disc. (A)-(A’’’)** Confocal scan on the level of the disc proper. Cells of the *pnr* lineage contribute to the dorsal disc margin (white arrowheads). A few cells of the *pnr*-lineage also contributed to the dorsal-posterior developing retinal field (open arrowheads). *pnr*>GFP expression can be detected in the margin cells (white arrowheads) posterior to the ocelli anlagen (grey arrowheads). **(B)-(B’’’)** Confocal scan on the level of the peripodial epithelium. Cells of the *pnr* lineage covered most of the dorsal part of the peripodial epithelium (white arrow), while *pnr* expression is restricted to the dorsal most region (grey arrow) in late L3 larval stages.

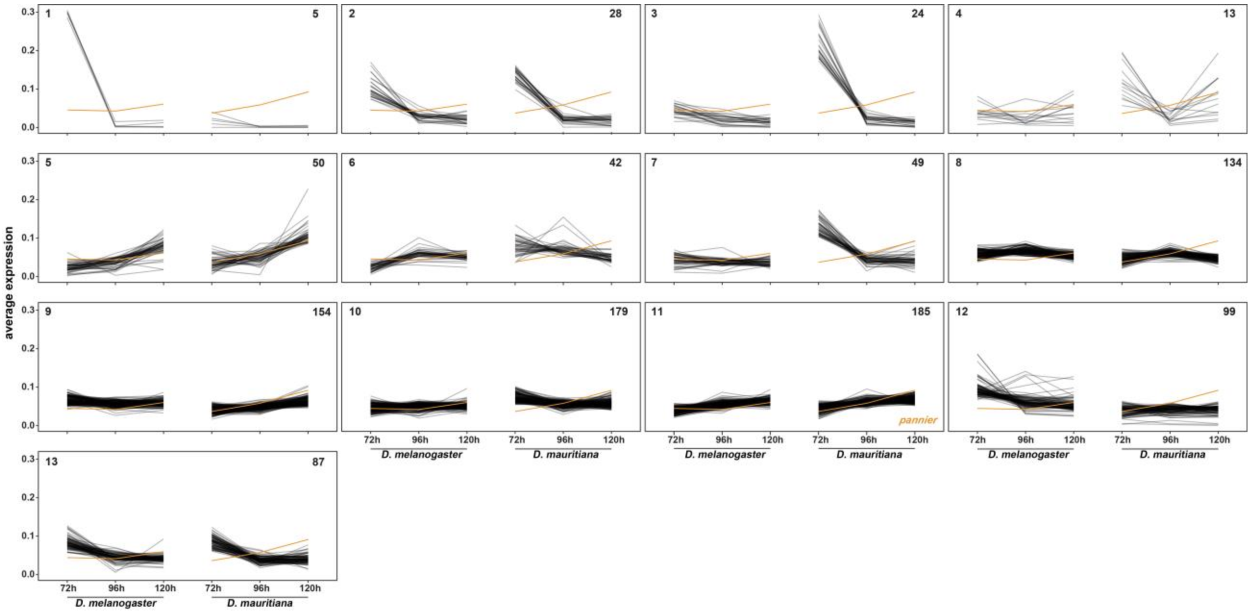


**S8 Fig. Clustering of differentially expressed putative Pnr target genes based on expression dynamics.** The number of genes in each cluster is provided in the top right corner. Among the 13 obtained clusters, *pnr* itself is present in cluster 11. The expression profile of *pnr* is provided in orange in each cluster for comparison. While other genes in cluster 11, as well as genes in clusters 5, 9 and 10 showed the same expression dynamics as *pnr*, genes in other clusters showed the exact opposite trend. For instance, the Pnr target genes in cluster 3 were highly expressed at 72 h AEL in *D. mauritiana*, while *pnr* itself showed a relatively low expression. The expression of the same target genes decreased at 120 h AEL with *pnr* expression increasing at the same time. This contrasting expression profile strongly suggests that those target genes may be repressed by Pnr action. In contrast, genes in clusters that show the same dynamics as *pnr* may be positively regulated by Pnr.

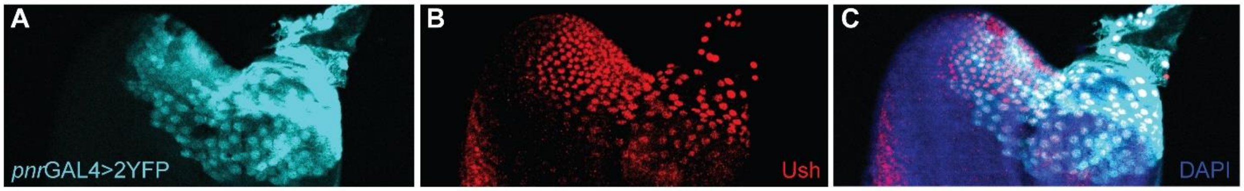


**S9 Fig. Co-expression of *pnr* and Ush in the peripodial membrane of the eye-antennal disc in *D. melanogaster*. (A)** *pnr* expression visualized by *pnr*GAL4>2YFP reporter line. **(B)** Ush protein localization detected with an α-Ush antibody. **(C)** Overlay of **(A)** and **(B)**. Posterior to the left and dorsal up.

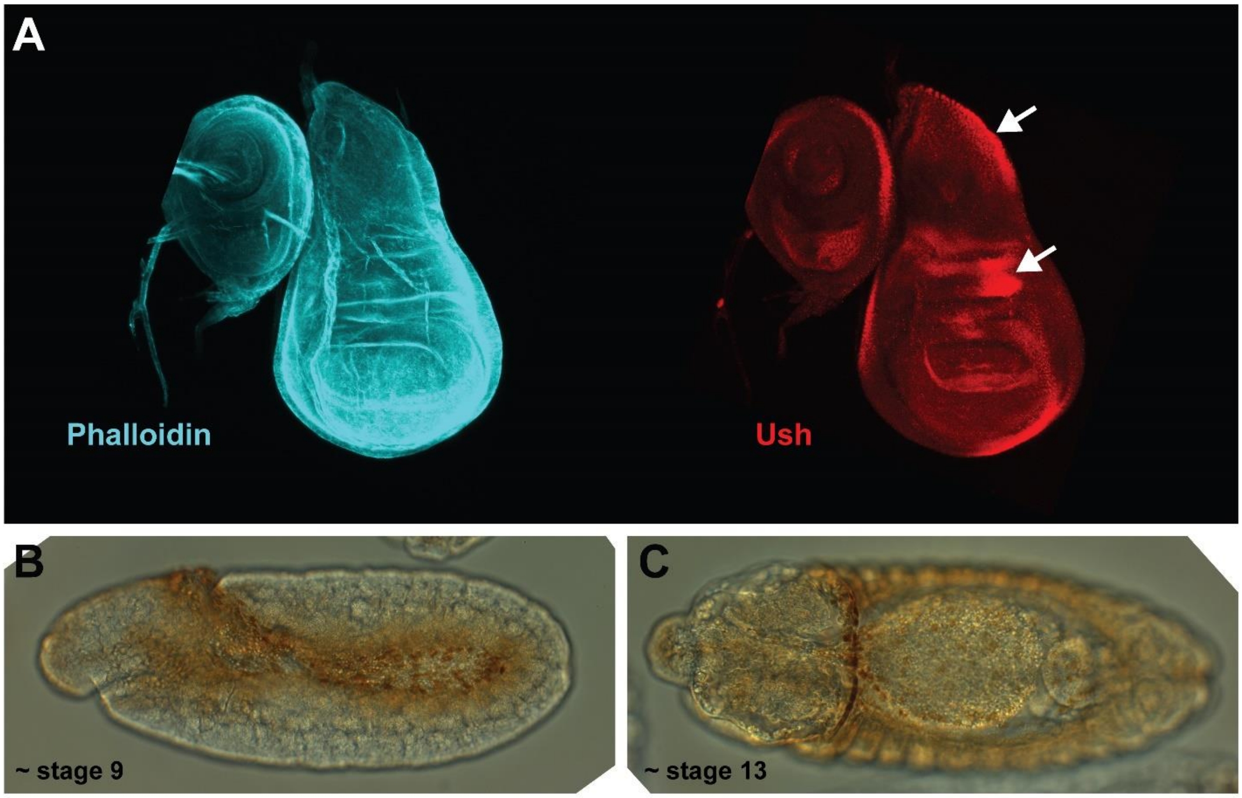


**S10 Fig. Expression of Ush in the wing imaginal disc and in the embryo. (A)** Ush protein location in the developing *Drosophila* wing disc, detected with the newly generated α-Ush antibody. The regions where Ush were detected in this tissue (white arrows) are reminiscent of the regions, where *ush* mRNA can be detected using *in-situ* hybridization as shown in [4]. **(B)** Ush protein location in a developing *Drosophila*

embryo at about stage 9, detected with the α-Ush antibody. Ush can be detected in the dorsal ectoderm and the head mesoderm, where also *ush* mRNA was detected, as shown in [5–8]. **(C)** Ush protein location in a developing *Drosophila* embryo around ~stage 13 in the leading cells of the future dorsal closure (also compare to *ush* mRNA in embryos stage 13-16 in [6–8]), detected with the α-Ush antibody.

**S1 Table. Number of differentially expressed genes (DEGs).**

| stage | upregulated in *D. melanogaster* | upregulated in *D. mauritiana* | total number of DEGs |
|---|---|---|---|
| **72h AEL** | 3,244 | 3,439 | 6,683 |
| **96h AEL** | 1,622 | 1,638 | 3,260 |
| **120h AEL** | 1,130 | 1,250 | 2,380 |

**S2 Table. GO enrichment and iCis-Target analyses for clusters of differentially expressed genes.**

See Excel file S2 Table

**S3 Table. List of putative Pnr target genes.**

See Excel file S3 Table

### Extended Material and Methods

#### Generation of the transcriptomic dataset

Flies from the following strains were raised at 25°C at a 12:12 dark:light cycle for at least two generations and their eggs were collected on agar plates for one hour: *D. melanogaster* (OreR), *D. mauritiana* (TAM16). 30 L1 larva were collected in vials and developing eye-antennal discs were dissected at 72h AEL (120–130 discs; male and female), 96h AEL (80–90 discs; female) and 120h AEL (40-50 discs; female) and stored in RNALater (Quiage, Venlo, Netherlands). For each species and stage 3 biological replicates were dissected. Total RNA was isolated using the Trizol (Invitrogen, Thermo Fisher Scientific, Waltham, Massachusetts, USA) method according to the manufacturer's recommendations and the samples were DNAseI (Sigma, St. Louis, Missouri, USA) treated in order to remove DNA contamination. RNA quality was determined using the Agilent 2100 Bioanalyzer (Agilent Technologies, Santa Clara, CA, USA) microfluidic electrophoresis. Only samples with comparable RNA integrity were selected for sequencing. Library preparation for RNA-seq was performed using the TruSeq RNA Sample Preparation Kit (Illumina, catalog ID RS-122-2002) starting from 500 ng of total RNA. Accurate quantitation of cDNA libraries was performed using the QuantiFluordsDNA System (Promega, Madison, Wisconsin, USA). The size range of final cDNA libraries was determined applying the DNA 1000 chip on the Bioanalyzer 2100 from Agilent (280 bp). cDNA libraries were amplified and sequenced (50 bp single-end reads) using cBot and HiSeq 2000 (Illumina). Sequence images were transformed to bcl files using the software BaseCaller (Illumina). The bcl files were demultiplexed to fastq files with CASAVA (version 1.8.2)

##### Mapping

The reads were mapped against strain-specific transcriptomes (*D. melanogaster* and *D. mauritiana*), including CDS and UTR [9] using Bowtie2 v. 2.3.4.1 [10] with the following parameters: -very-sensitive-local -N1. Samtools version 1.9 [11,12] was used to further process the reads and count the reads mapped to each transcript (idxstats).

##### Differential expression analysis and data visualization

The PCA plot is based on the regularized log (rlog) transformation from the DESeq2 package (DESeq2_1.22.2 [13]; R version 3.5.2).

DeSeq2 was used to perform a pairwise differential expression analysis between the two species at each time point using the apeglm option as shrinkage estimator [14] (*D. melanogaster* 72h vs. *D. mauritiana* 72h, *D. melanogaster* 96h vs. *D. mauritiana* 96h and *D. melanogaster* 120h vs. *D. mauritiana* 120h). We used the online tool Metascape [15] to perform GO enrichment analysis for each time point. We combined the read counts of all genes that were significantly differentially expressed (log2FC > 0 | log2FC < 0 and padj < 0.05) between the two species in at least one stage (8,350 genes in total). We then clustered them according to their expression dynamics (i.e. we treated the combined data from both species as a time-series) using the coseq package (version 1.6.1) [16,17] with the following parameters: K=2:25, transformation="arcsin", norm="TMM", model="Normal". We searched for potential upstream factors 5 kb upstream of the transcription start site, 5'UTR regions and $1^{st}$ introns using the i-cisTarget tool [18,19] keeping the default parameters: Minimum fraction of overlap: 0.4, NES: 3.0, ROC threshold for AUC calculation: 0.01. S2 Table contains enriched unique motifs with NES > 4.0, we excluded non-*Drosophila* motifs, and added only the ones which could be found on Flybase [20]. To check which of these potential upstream regulators are differentially expressed, we removed all duplicates and overlapped the list with all significantly differentially expressed genes.

Metascape was used to analyse differential enrichment of GO terms for each pairwise comparison.

#### Generation of the ATACseq dataset

For the generation of ATAC-seq datasets we followed a previously established protocol [21]. Developing eye-antennal discs of *D. melanogaster* were dissected in ice-cold PBS at 72h, 96h and 120h AEL. PBS was removed and exchanged for 50 µl lysis buffer (10 mM Tris-HCl (pH = 7.4); 10 mM NaCl; 3 mM MgCl2; 0.1 % IGEPAL). The mixture was pipetted several times up and down to lyse the cells and then split into micro centrifuge tubes. Centrifugation for 10 min at 500 g and 4°C. The cell number was assessed in one of the samples and between 50,000 and 80,000 nuclei were used in subsequent steps. The supernatant was removed, and the pellet(s) were dissolved in 47.5 µl 1X tagmentation buffer (20 mM Tris-CH3COOH (pH = 7.6); 10 mM MgCl2; 20 % (vol/vol) dimethylformamide) with 2.5 µl Tn5 Transposase and then incubated for 30 min at 37°C. For purification we used the QIAGEN MinElute Kit and eluted in 10 µl Elution Buffer (10 mM Tris, pH = 8). The PCR amplification was done as follows: 10 µl tagmented chromatin,

10 µl $H_2O$, 2.5 µl Nextera PCR primer 1*, 2.5 µl Nextera PCR primer 2**, 25 µl NEBNext High-Fidelity 2X PCR Master Mix (Cat #M0541).

* AATGATACGGCGACCACCGAGATCTACACTCGTCGGCAGCGTCAGATGTG

** Ad2.2_CGTACTAG
CAAGCAGAAGACGGCATACGAGATCTAGTACGGTCTCGTGGGCTCGGAGATGT
Ad2.3_AGGCAGAA
CAAGCAGAAGACGGCATACGAGATTTCTGCCTGTCTCGTGGGCTCGGAGATGT
Ad2.4_TCCTGAGC
CAAGCAGAAGACGGCATACGAGATGCTCAGGAGTCTCGTGGGCTCGGAGATGT
Ad2.5_GGACTCCT
CAAGCAGAAGACGGCATACGAGATAGGAGTCCGTCTCGTGGGCTCGGAGATGT
Ad2.6_TAGGCATG
CAAGCAGAAGACGGCATACGAGATCATGCCTAGTCTCGTGGGCTCGGAGATGT
Ad2.7_CTCTCTAC
CAAGCAGAAGACGGCATACGAGATGTAGAGAGGTCTCGTGGGCTCGGAGATGT

We used the following program: 1) 72 °C for 5 min, 2) 98 °C for 30 sec, 3) 98 °C for 10 sec, 4) 63 °C at 30 sec, 5) 72 °C for 1 min, step 3) -5) were repeated 13 times. This was followed by another 2x purification step with the QIAGEN MinElute Kit: elution in 2 X 10 µl Elution Buffer (10 mM Tris, pH = 8).

#### Bioinformatic processing of the ATACseq data

We performed quality checks of the sequenced reads using FASTQC (https://www.bioinformatics.babraham.ac.uk/projects/fastqc/). The reads were trimmed using Trimmomatic (version 0.36) [22] with the parameters slidingwindow 4:15 and minlen 30. Trimmed reads were mapped to the *D. melanogaster* genome (version 6.13) (after discarding the mitochondrial genome) using Bowtie2 (version 2.3.4.3), with the commands: --no-unal and -X2000. Samtools version 1.9 was subsequently used to convert the sam files to bam files, and to sort and index bam files. We removed duplicates using PICARD (version 2.1.1., http://broadinstitute.github.io/picard/) with default parameters and converted the resulted bam files to bed files. Reads were then centred as described in [21]. We used MACS2 (version 2.1.2) [23] with the following commands: -g dm --nomodel --shift -100 --extsize 200 -q 0.01 –bdg to call significant peaks. We used the Integrated Genome Browser (IGB) [24] to visualize the read depth and peaks. Peaks were annotated to the closest gene using the annotatePeaks.pl program from the HOMER software package (v4.8.3) [25] using dm6 as genomic input.

#### Definition of a Pnr target gene list

As a basis for the high confidence list of putative Pnr target genes a Chip-chip dataset was used (downloaded on 1st of July, 2015 from http://furlonglab.embl.de/data/download), which comprises ChIP-chip experiments in the *Drosophila* embryo with several transcription factors, including Pnr at two time points (4-6h AEL and 6-8h AEL) [26]. All Pnr-binding regions from both time points were selected with a Tile-Map score of <5.5. and where the distance of the centre of the peak to the transcription start site was -1000 bp and +1000 bp. We performed a *de novo* motif search in these peak regions, to define the Pnr binding motif. We used this motif to screen a set of combined and unique ATAC-seq peaks from three time-points for potential Pnr binding sites in open chromatin regions of the developing eye-antennal disc, which resulted in 1,335 unique peaks in total. These peaks were annotated to 1,241 genes, of which 1,060 (S3 Table) are expressed in the eye antennal disc (>= 10 reads in at least one of the three stages). We overlapped these genes with all genes that were significantly differentially expressed in at least one of the three stages (8,350 genes). To understand the expression dynamics of potential Pnr target genes in relation to Pnr itself, we performed hierarchical clustering according to their expression dynamics using coseq with the following parameters: K=2:25, transformation="arcsin", norm="TMM", model="kmeans".

We downloaded all known direct (TF-gene, 157,462 interactions) or genetic interactions (14,241 interactions) from the DroID database [27,28] (in total 171,703 interactions). We extracted from this database all interactions of Pnr with other genes and found 181 target genes of Pnr. 26 of these were present our Pnr target gene list and 17 were differentially expressed. We used Cytoscape [29] to visualize the interaction between these target genes and potential upstream regulators found in the database.

#### Antibody generation and immunohistology

We generated polyclonal antibodies against Pnr [26] and Ush [30] based on previous knowledge against the following peptide sequences (Proteintech, Rosemont, IL, USA):

Pnr_B (125-294):
TPLWRRDGTGHYLCNACGLYHKMNGMNRPLIKPSKRLVSATATRRMGLCCTNCGTRT
TTLWRRNNDGEPVCNACGLYYKLHGVNRPLAMRKDGIQTRKRKPKKTGSGSAVGAGT
GSGTGSTLEAIKECKEEHDLKPSLSLERHSLSKLHTDMKSGTSSSSTLMGHHSAQQ

Pnr_B (206-336):

GVNRPLAMRKDGIQTRKRKPKKTGSGSAVGAGTGSGTGSTLEAIKECKEEHDLKPSLSL
ERHSLSKLHTDMKSGTSSSSTLMGHHSAQQQQQQQQQQQQQQQQQQQSAHQQCFPL
YGQTTTQQHQQHGH

Ush (231-250):

CSHRIKDTDEAGSDKSGAGG

Ush (1174-1191):

VGGHGQQKNKENLQEAAI

Before usage, both antibodies were preabsorbed overnight in a mix of *Drosophila* embryos on 4°C.

Developing eye-antennal discs were dissected and fixed for 30 min in 4% paraformaldehyde (PFA). For this purpose, PFA was dissolved in $ddH_2O$ and 1M NaOH by boiling, diluted with 10x Phosphate Buffered Saline (PFA) and the pH was adjusted to 7.2 using NaOH. The discs were then washed 3 times in 0.03% PBT (1x PBS, 0.03% Triton X-100) before blocking in 5% normal goat serum for 30 min. Incubation with the primary antibody was done for 90 min, before 3 additional washing steps with PBT and one round of blocking for 30 min. The discs were then incubated with the secondary antibody overnight on a rocking plate at 4°C. Phalloidin-488 (1:100) was added. After 3 washing steps with PBT, the discs were incubated with DAPI (1:1000) for 10 min, followed by one washing step with PBT and one washing step with PBS. Subsequently, the discs were mounted in mounting medium (80% glycerol + 4% n-propyl-galate) and kept at least one night on 4°C prior to imaging.

We confirmed the specificity of the Ush antibody by recapitulating known Ush expression domains in the wing imaginal disc (S10A Fig) and during embryonic development (S10B and S10C Fig). For test stainings in embryos we collected embryos for several hours on apple agar plates, removed the chorion with 50% bleach (DanKlorix) and rinsed them 3 times with 0.03% PBT (Phosphate buffered saline 1%, Triton X-100). We fixed the embryos with heptane and 2% formaldehyde for 20min and washed with MeOH, followed by washing steps with PBT. The embryos were then blocked in 3% BSA for one hour, followed by incubation with the primary antibody overnight. After two washing steps with PBT, we added HRP-coupled secondary antibody for 90 min. After three washing steps with PBT we performed a DAB (3’-

3diaminobenzidine) staining. The embryos were then washed again 2 times in PBT and mounted in glycerol.

Concentrations of antibodies used: Anti-Pnr (rabbit): 1:200 (Proteintech, Rosemont, IL, USA), Anti-Ush (rabbit): 1:2000 (Proteintech, Rosemont, IL, USA), Anti-GFP (chicken): 1:1000 (Abcam # 13970, kindly provided by Dr. Marita Büscher); Concentrations Secondary antibodies: Anti-Chicken-488 (Invitrogen Cat# A-11039, kindly provided by Dr. Marita Büscher); Anti-Rabbit-Cy3 (donkey): 1:500 (Jackson ImmunoResearch #711-165-152, kindly provided by Kolja N. Eckermann); Anti-rabbit-HRP-coupled (1:1000) (Jackson ImmunoResearch #111-035-003, kindly provided by Dr. Marita Büscher).

Pictures of eye-antennal discs upon antibody staining were taken using a Zeiss LSM 710 confocal microscope. Antibody stainings were visualized and processed with Fiji software. Vertical sections of the confocal pictures were generated using the Volume Viewer plugin using the following parameters: Display Mode: Slice and Boarders, Interpolation: Nearest Neighbour, Transfer Function: Fire LUT.

#### Geometric Morphometrics

We imaged dorsal heads of wildtype specimens from each parental line, as well as the offspring of the respective *D. melanogaster* crosses using a Leica M205 FA stereo microscope. We placed 64 landmarks on pictures of these dorsal heads using the tpsDig2 software [31]. We then defined 23 fixed landmarks and 41 sliding landmarks using tpsUtil [31]. tpsRelw32 [31] was used to calculate the consensus (i.e. Procrustes superimposition), partial warps and relative warps. Using MorphoJ [32] for visualization, we performed Procrustes Fit and generated a covariance matrix. To analyse differences in dorsal head shapes in wildtype *D. melanogaster* and *D. mauritiana*, we performed canonical variate analysis (CVA) using MorphoJ. We further performed a principal component analysis (PCA) to analyse difference in head shapes upon knock-down or up-regulation of *pnr*.

#### Quantitative analysis of eye-antennal disc development

Eye-antennal discs were dissected as described above ("Generation of the transcriptomic dataset"). The discs were fixated, incubated with Phalloidin-488 (1:100) and imaged as described above ("Antibody generation and immunohistology"). The linear measurement tool implemented in the Fiji software was used to measure distances as implicated in Fig 1E and 1F. The area of the antennal field was measured using the area measurement tool as implemented in Fiji (Fig 1D). The number of future ommatidia rows was counted from the optic stalk to the morphogenetic furrow along the equator of the retinal field (Fig 1G).

#### Overexpression/Knock-down of *pnr* and *ush*

To overexpress or knock-down *pnr* and *ush*, the following fly lines were used:

*D. melanogaster* (Oregon R) and *D. mauritiana* (TAM 16) (both kindly provided by Alistair McGregor, Oxford Brookes University), *pnr* GAL4/TM6B (kindly provided by Marc Haenlin), *pnr*-GAL4>UAS2YFP/TM6B (kindly provided by Marc Haenlin, CBI Toulouse), *ush*-GAL4 26662 (y[1] w[*]; P{w[+mW.hs]=GawB}*ush*[MD751]), Bloomington; UAS-*ush*14IIA/CyO (kindly provided by Marc Haenlin, CBI Toulouse); *ush*-RNAi 3622 (y[1] v[1]; P{y[+t7.7] v[+t1.8]=TRiP.HM05193}attP2/TM3, Sb[1]), Bloomington; UAS-*pnr* (w; UAS-*pnr*/CyO; TM2/TM6B) (kindly provided by Fernando Casares); *pnr*-RNAi (VT101522/KK, #108962 VDRC Stock Center and VT6224/GD, #1511 VDRC Stock Center); *oc*-Gal4/CyO (kindly provided by Fernando Casares); UAS-Stinger-GFP (nGFP) (kindly provided by Dr. Gerd Vorbrüggen).

All crosses were performed at 25°C and at a constant 12h:12h light:dark cycle. Since Pnr and Ush are crucial during embryonic development, we chose combinations of GAL4/UAS lines that resulted in a phenotype, but were not lethal during embryonic, larval or pupal stages. We used a set of GAL4 lines representing a range from weak to strong drivers and in the case of *pnr*-RNAi we used two independent RNAi lines.

#### Pnr expression and lineage tracing

The *pnr-GAL4* line *pnrMD237* [33], recombined with UAS-GFP, was used to follow *pnr* expression in imaginal and pupal eye-antennal discs. The adult *pnr* expression domain in adult heads was monitored in *y; pnr-GAL4/UAS-y+* flies [33] as the cuticle region with y-rescued pigmentation. To follow the *pnr-GAL4* lineage, *pnr-GA4*, UAS-GFP flies were crossed to UAS-flipase; *act5c>stop< nuc-lacZ* flies [34]. In the discs of the progeny, actual *pnr* expression was visualized with anti-GFP and its lineage with anti β-galactosidase.

##### Immunostaining and imaging.

Third instar or pupal discs were processed as in [35]. Primary antibodies were chicken anti-GFP (1:500; ab13970, Abcam), rabbit anti-β-galactosidase (1:1000; Cappel) and mouse anti-Eya (1:500; 10H6, Developmental Studies Hybridoma Bank, Iowa University). Secondary antibodies at 1:400 were from Molecular Probes. Imaging was carried out on a Leica SPE confocal setup (ALMI, CABD).

##### Adult head cuticle preparation.

Dissected adult heads in PBS were mounted in Hoyer's medium: Lactic Acid (50:50) as in [35].

##### Ommatidia Counting

To estimate ommatidia number of single fly eyes, we took pictures of one eye per fly (50 stacks/eye) using a Leica M205 FA stereo microscope and an external light source, which resulted in reflection of light by each ommatidium. We used FIJI to perform Z-projections using maximum intensity and then transformed each picture to make sure that the single reflection of each ommatidium is represented by a black dot (i.e. increase of contrast, transform to grey scale and invert photo). We then used the ITCN cell counter tool (with the following parameters: Width: 7, Minimum Distance: 17, Threshold: 1.5; https://bioimage.ucsb.edu/search/node/itcn) to estimate the number of black dots, i.e. the number ommatidia. Statistical analysis of ommatidia number was done using a one-way ANOVA and pair-wise comparisons were calculated using Tukey HSD test.
